# Personalized Whole-Brain Models of Seizure Propagation

**DOI:** 10.1101/2025.05.15.654238

**Authors:** Edmundo Lopez-Sola, Borja Mercadal, Èlia Lleal-Custey, Ricardo Salvador, Roser Sanchez-Todo, Fabrice Wendling, Fabrice Bartolomei, Giulio Ruffini

## Abstract

*Objective:* Computational modeling has recently emerged as a powerful tool to better understand seizure dynamics and guide new treatment strategies. This work aims to develop and personalize whole-brain computational models in epilepsy using multimodal clinical data to simulate and evaluate individualized therapeutic strategies. *Approach:* We present a computational framework that constructs patient-specific whole-brain models of seizure propagation by integrating SEEG, MRI, and diffusion MRI data. The pipeline uses neural mass models for each node in the network, simulating whole-brain dynamics. Model personalization involves adjusting global and local parameters representing the excitability of individual brain areas, using an evolutionary algorithm that aims to maximize the correlation between empirical and synthetic functional connectivity matrices derived from SEEG data. *Main results:* The resulting personalized models successfully reproduce individual seizure propagation patterns and can be used to simulate therapeutic interventions like surgery, stimulation, or pharmacological interventions within a unified physiological framework. Notably, model predictions reveal distinct patient-specific responses across interventions, including variable sensitivity to different pharmacological agents and identification of critical regions whose removal or modulation reduced seizure spread. *Significance:* This framework provides a mechanistic, interpretable approach to simulate and compare individualized treatment strategies. By integrating multimodal data into a unified whole-brain model, it has the potential to improve clinical decision-making in epilepsy by identifying accessible and functionally relevant targets.

## 1 Introduction

Epilepsy is one of the most common neurological disorders worldwide, affecting approximately 50 million people [1]. It is characterized by an enduring predisposition to epileptic seizures and by cognitive, psychological, and social comorbidities that impose a significant burden on individuals and healthcare systems. Among those affected, a substantial subset of patients suffers from drug-resistant epilepsy (DRE), a form that proves resistant to conventional treatments such as antiseizure drugs. Approximately 30% of epilepsy patients with DRE continue to experience seizures despite treatment [2]. For these patients, epilepsy remains a debilitating disease with few effective solutions, often leading to a lower quality of life due to the ongoing unpredictability of seizure events and associated risks such as injury and deterioration of mental health.

To better understand the mechanisms underlying seizures and explore new treatment strategies, computational modeling has emerged as a valuable tool in epilepsy research [3–5]. These models have been applied in various contexts, including i) the identification of the epileptogenic (EZ) and propagation zones (PZ) [6–10], ii) the prediction of surgical outcomes [11–19], iii) the interpretation of intracranial and scalp EEG [20–24], and iv) the design of neurostimulation strategies [25, 26]. Depending on the specific goal of the study, different modeling approaches have been adopted, each with distinct trade-offs between physiological interpretability and phenomenological accuracy.

For instance, phenomenological models (such as the Epileptor [6]) have been widely used to replicate seizure dynamics observed in electrophysiological recordings, particularly for applications related to surgical planning and the identification of regions generating and propagating seizures [4, 7, 17]. These models are designed to capture key features of seizure transitions with relatively few parameters, allowing for efficient personalization and large-scale simulations. However, their abstraction from the underlying physiology limits their use in studies requiring neurobiologically-based interpretations, such as the simulation of pharmacological effects or electrical stimulation.

In contrast, neural mass models (NMMs) operate at the mesoscale and offer a richer neurophysiological interpretation [3]. These models describe the average activity of interconnected neuronal populations and include parameters related to synaptic dynamics, membrane potentials, and firing rate transformations. Due to their structure, NMMs are particularly suited for simulating interventions that depend on physiological detail, such as neuromodulation or pharmacology, and have also been used to investigate the propagation of seizures and evaluate surgical strategies [8, 12, 15]. Other studies have explored even simpler frameworks, such as bistable models[13, 27, 28], theta models [9, 16], or Wilson-Cowan formulations [25], which offer a compromise between computational efficiency and biological realism. Building on previous work on neural mass modeling in epilepsy [20, 21], we have recently developed a novel neural mass that generates realistic transitions from pre-ictal to ictal state [29]. We have also introduced a framework for the integration of NMMs in a layered physical model of a cortical column [30], as well as a physical model for the simulation of SEEG signals from neural masses [31]. Those developments are crucial for the design of the model personalization pipeline presented in the current work.

Although computational models offer insight into the mechanisms of the generation and spread of seizures, their potential extends beyond understanding the pathology. One of their most promising applications is in the simulation and optimization of therapeutic strategies. Among these, brain stimulation and neuromodulation have emerged as key candidates for intervention, particularly in patients who do not respond to conventional pharmacological treatments [32, 33]. Transcranial Direct Current Stimulation (tDCS) has attracted attention for its safety profile, ease of use, and ability to modulate neuronal excitability with minimal discomfort (VNS) [34], in contrast to more invasive techniques such as Deep Brain Stimulation (DBS) or Vagus Nerve Stimulation (VNS). tDCS operates through the application of a low-intensity, constant current delivered directly to the brain areas via electrodes positioned on the scalp [35]. In previous work, we have demonstrated the efficacy of tDCS in refractory focal epilepsy by targeting the epileptogenic focus with multichannel montages, optimized using personalized biophysical head models of the patients, which included anatomical data and simulation of the head tissues and conductivities [26, 36, 37].

Building on this, we aim to leverage personalized, whole-brain computational models that include not only biophysical but also physiological information about the patient’s brain, such as electroencephalography (EEG) or stereo-EEG (SEEG). Indeed, whole-brain computational models represent a transformative approach to epilepsy treatment, providing *in silico* replication of a patient’s neural activity and mechanistic insight into the network dynamics and pathophysiological processes underlying epileptic seizures. Our vision is that such models can be effectively used to simulate and predict the outcomes of various treatment scenarios, including surgery and stimulation therapies, improving the results achieved to date using personalized biophysical models.

In this paper, we present a method for personalizing whole-brain computational models in epilepsy by integrating SEEG, anatomical MRI, and diffusion MRI (dMRI) data. Unlike many existing approaches that aim to localize the seizure onset zone [6, 9, 16], our focus is not on identifying the epileptogenic focus itself. Instead, we rely on the clinician’s identification of an epileptogenic network and attempt to construct a mechanistic model that captures how seizures propagate from this network to the rest of the brain. To do so, we incorporate both structural connectivity, derived from patient-specific dMRI, and functional information extracted from SEEG recordings using amplitude envelope correlations. This integration allows us to personalize the excitability of different brain areas and capture early seizure propagation patterns in a data-driven manner.

The motivation for this approach stems from a clinical challenge: interventions such as surgery or non-invasive stimulation often target the presumed seizure onset zone but do not always lead to improved outcomes [38]. This suggests that seizure spread, and not just seizure initiation, plays a critical role in treatment response. By modeling the dynamics of seizure propagation, we aim to identify additional targets that contribute to seizure spread and may be amenable to therapeutic modulation. Ultimately, our goal is to provide personalized models that can inform interventions aimed at reducing both seizure frequency and spatial extent. This is particularly relevant in cases where the seizure onset zone is located in deep brain structures that are difficult to reach with surgery or non-invasive techniques like tDCS, making it essential to consider propagation targets that are more accessible.

In the following sections, we detail the modeling pipeline and demonstrate how it can reproduce the empirical seizure propagation observed in SEEG data. We also show that the resulting models can simulate and compare the effects of various interventions on seizure dynamics in a patient-specific manner.

## 2 Methods

### 2.1 Overview

First, we present an overview of the full personalization pipeline (see Figure 1). The inputs to the pipeline are the SEEG data during a seizure, the subject’s MRI and dMRI, the locations of the SEEG contacts in the Virtual Epileptic Patient (VEP) parcellation [39], and the clinician’s hypothesis of the EZ location. The processed data is used to build the patient’s whole-brain model, and the model parameters are personalized using an evolutionary algorithm that maximizes the correlation between the empirical and synthetic functional connectivity (FC) matrices. The output is the set of optimized model parameters, which define the personalized whole-brain model of the patient. After model personalization, the validity of the model can be assessed by comparing the seizure propagation pattern in the data and the model. The model can then be used to predict the effects of therapeutic interventions.

**Figure 1:**
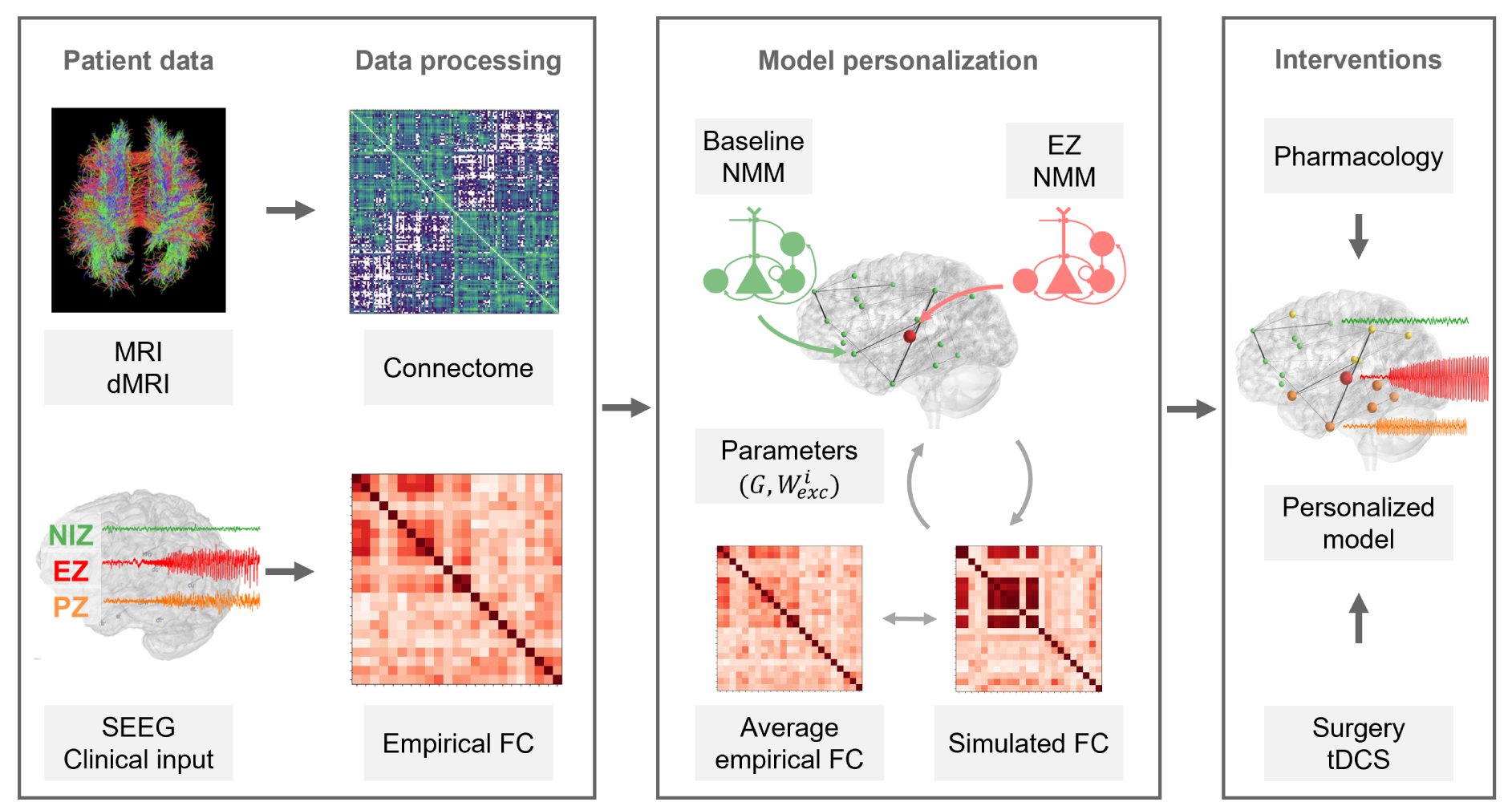
Overview of the full personalization pipeline. Patient-specific data, including structural MRI, dMRI, and SEEG, are processed to extract the structural connectome and empirical functional connectivity. Regions are classified by an expert clinician according to their epileptogenicity. A whole-brain model composed of neural mass models is personalized by tuning global and local excitability parameters (*G*, *W*_exc_), to match simulated FC with empirical FC. The resulting model is used to simulate therapeutic interventions such as pharmacology or surgery. Abbreviations: MRI = magnetic resonance imaging; dMRI = diffusion MRI; SEEG = stereoelectroencephalography; NMM = neural mass model; FC = functional connectivity; EZ = epileptogenic zone; PZ = propagation zone; NIZ = non-involved zone; tDCS = transcranial direct current stimulation.

### 2.2 Patient data and data processing

#### 2.2.1 Clinical data

The patients included in this study required SEEG as part of their routine clinical care. Clinical information about the patients can be found in Table 1. In all cases, multiple intracerebral electrodes were placed in different brain regions of the patient based on the clinician’s hypotheses about the location of the epileptogenic zone.

**Table 1:** Clinical characteristics of the patients included in the study.

| Patient ID | 1 | 2 | 3 | 4 | 5 | 6 |
| --- | --- | --- | --- | --- | --- | --- |
| Gender | F | F | M | M | F | M |
| Age at epilepsy onset (y) | 14 | 9 | 11 | 12 | 34 | 19 |
| Epilepsy duration (y) | 7 | 14 | 4 | 32 | 10 | 16 |
| Epilepsy type | Parietal | Occipital | Temporal | Frontal | Temporo-insular | Temporal |
| Surgical procedure | SR | SR | TC, SR | TC, SR | SR | LiTT |
| Surgical outcome (Engel class) | I | II | I | I | I | I |
| MRI | R parietal DNET | Normal | Normal | Normal | Normal | Normal |
| Histopathology | DNET | FCD1 | FCD2 | FCD2 | Non specific | Not available |
| Side | R | L | L | R | R | L |
| Seizures recorded | 3 | 3 | 2 | 3 | 1 | 1 |

Abbreviations: DNET, dysembryoplastic neuroepithelial tumor; FCD, focal cortical dysplasia; L, left; R, right; SR, surgical resection; TC, thermocoagulations; LiTT, Laser interstitial thermal therapy.

The data used in this study is part of the Galvani project (ERC Synergy Grant 855109). Patients 4 and 5 participated in a pilot study conducted within the framework of this project to evaluate the impact of tDCS on SEEG recordings [40], and Patient 6 was included in a separate pilot study also related to the project to assess the efficacy of tDCS in DRE [26]. For this work, we selected all patients for whom the complete dataset was available, including SEEG recordings during seizures, diffusion MRI (dMRI), and anatomical MRI.

#### 2.2.2 SEEG data and contact selection

SEEG recordings were obtained using intracerebral multiple-contact electrodes with a 154-channel Deltamed system. Contacts were classified by the clinician according to their epileptogenicity into three distinct networks [41]:

- Epileptogenic Zone (EZ): the network responsible for the generation of seizures.
- Propagation Zone (PZ): a less epileptogenic network that propagates seizures but does not initiate them.
- Non-involved zone (NIZ): areas of the cortex not belonging to the EZ or PZ.

The contact locations in the VEP parcellation atlas were obtained using the GARDEL software [42]. The VEP atlas has been shown to provide an effective balance between anatomical resolution and computational feasibility for modeling SEEG data [18, 39, 43]. Bipolar derivations were obtained from neighbor contact subtraction for each electrode. A parcel was assigned to each bipolar pair based on the individual contact locations. In the case where the contacts used for bipolar derivation were assigned to different parcels, we chose the parcel of the first contact in alphabetical/numerical order. Bipolar pairs where neither contact belonged to the grey matter were discarded.

The empirical FC computation for the modeling pipeline required a single SEEG bipolar signal for each parcel (see Section 2.2.4). Since several contacts could be assigned to the same parcel, we defined a method for selecting one bipolar signal for each parcel sampled by SEEG, inspired by the method used in Olmi et al. [17] to identify the propagation zone:

1. Filter each bipolar signal between 1-50 Hz using a Butterworth band-pass filter.
2. Compute the *energy ratio* of each contact pair as the ratio between the signal energy during the seizure period (marked by the clinician) and the baseline signal energy (one minute before and one minute after the seizure).
3. Identify which contacts belong to the EZ or PZ according to the clinician.
4. Select one contact pair per parcel by choosing the most epileptogenic pair (EZ before PZ before NIZ), and in the cases where several pairs had the same epileptogenicity level, choose the contact pair with the highest mean energy ratio (average over seizures).

This method ensured that we included in the FC computation the maximum number of EZ and PZ contacts possible, and that the contacts selected were the ones with the highest energy in the seizure period relative to baseline. This change in energy is intended to capture the potential changes in amplitude due to seizure propagation, which is a key feature in the computation of the empirical FC (see Section 2.2.4).

#### 2.2.3 Structural connectivity

The individual structural connectivity of each patient was generated from diffusion MRI scans and T1-weighted images acquired on 3T MR scanners. Each MRI was segmented into 162 cortical regions according to the VEP atlas [39] using FreeSurfer [44] and custom Python scripts. This volumetric image served as the basis for processing dMRI data.

Diffusion MRI data were acquired using different acquisition protocols depending on the scanner model and imaging guidelines in place at the time of each patient’s screening. Although all protocols included single-shell acquisitions with b = 1000 s/mm^2^ and at least 64 diffusion directions, parameters such as echo spacing, readout time, and phase encoding directions varied between sessions. Some patients were scanned with both anterior-posterior (AP) and posterior-anterior (PA) phase encoding directions to enable distortion correction, while others had only a single direction available. More details about the acquisition parameters are provided in the Supplementary Information.

Preprocessing steps for diffusion MRI included denoising, artifact correction (e.g., eddy currents), and fiber orientation density estimation using constrained spherical deconvolution. Probabilistic tractography was then performed using MRtrix3 [45], generating 20 million streamlines constrained to start and terminate in gray matter parcels defined by the MRI parcellation. The resulting structural connectome matrix quantified the number of streamlines (fiber tracts) connecting each pair of parcels and was normalized by the total number of fibers divided by the number of parcels to generate a connectivity matrix for the subject.

#### 2.2.4 Empirical FC computation

Detecting the presence or absence of seizures in SEEG data is a complex problem, and several methods have been recently proposed for that purpose [41, 46]. Rather than trying to determine whether or not a given SEEG channel displays seizure, we have decided to use functional connectivity (FC) between signals to capture the pattern of seizure propagation in empirical data. Other studies have also used functional connectivity between SEEG signals for seizure onset localization or as markers of epileptogenicity [47, 48].

In the current pipeline, we computed the FC as the correlation between the amplitude envelopes of signal pairs [49]. This choice is based on the assumption that nodes involved in seizure propagation show pronounced amplitude increases that are temporally correlated across regions. In contrast, regions not participating in the propagation are expected to show irregular or low-amplitude fluctuations, resulting in quasi-random amplitude envelopes with low correlation to other signals. The use of amplitude envelopes for the fitting of model parameters has been employed in other studies of whole-brain model personalization in epilepsy [6]. It is worth mentioning that our FC computation method assumes that the SEEG electrodes capture a pronounced amplitude increase at seizure onset, which has some limitations that we discuss in Section 4.1.

##### Amplitude envelope computation

For the computation of the amplitude envelope correlation, we selected for each seizure a time window spanning from 10 seconds before the start of the clonic phase activity to 20 seconds after the start of the clonic phase, similarly to previous studies [47]. This window is intended to capture the early propagation of the seizure, which is the period relevant for the design of interventions aimed at reducing seizure spread (since targeting the earliest propagating nodes is expected to yield better outcomes in reducing seizure spread).

The initial estimation of the clonic phase onset was made by visual inspection of SEEG signals recorded from the EZ contacts specified by the clinician. We identified the point at which the signal began to show an increase in amplitude along with emerging rhythmic activity, consistent with typical clonic phase characteristics described in the literature [50]. To minimize variability introduced by manual selection, we explored a window from –10 to +10 seconds around the visually identified time point, in 1-second steps. For each candidate time point, we computed the corresponding FC matrix using the method described below and calculated the total node strength of EZ nodes as the sum of their connectivity values. The final clonic phase onset was defined as the time point that yielded the maximum EZ node strength, under the assumption that connectivity between EZ nodes peaks during early seizure propagation [51].

We first band-pass filtered each signal between 0.5 Hz and the Nyquist frequency (half the sampling rate), then computed the amplitude envelope of each signal as the absolute value of its Hilbert transform, and finally low-pass filtered the result below 1 Hz to smooth the envelope. The low-pass filtering of the amplitude envelope below 1 Hz helps emphasize slow fluctuations in signal power in the delta frequency band, which are more likely to reflect sustained, functionally relevant interactions between regions, rather than transient, high-frequency fluctuations or noise. We removed the first and last seconds of the signal to avoid the transients appearing after the Hilbert transform. The amplitude envelope was then normalized to its standard deviation and demeaned. This allowed us to capture amplitude increases in the SEEG signals independently of the absolute differences in signal intensity (see Figure 2).

**Figure 2:**
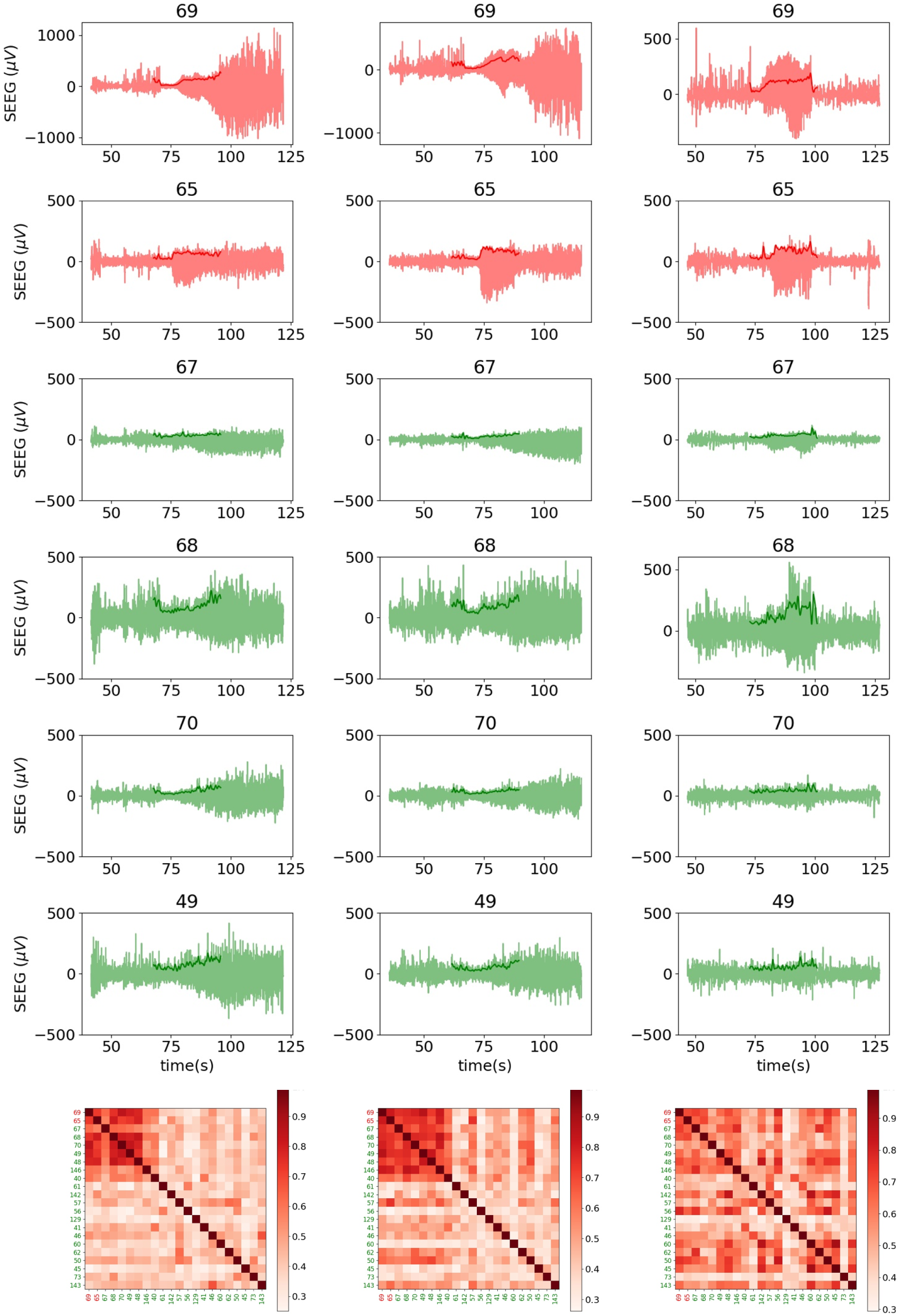
SEEG signals for the three seizures recorded in Patient 2 and corresponding empirical FC (bottom). Each column corresponds to a different seizure recording. The main EZ channels are shown (red), as well as four NIZ channels (green), are displayed, as well as their amplitude envelope (solid line above signals). Numbers correspond to the VEP parcel number [39] where the SEEG contacts are located. FC colorbars (bottom) represent PCC values between regional SEEG signals.

##### FC matrix computation

Finally, the FC of each pair of signals was computed as the maximum of the absolute value of the cross-correlation between the amplitude envelopes of the two signals. Using the maximum value allowed us to account for time delays, which are likely to occur due to delays in seizure propagation. Estimating and fitting these delays would make the personalization process significantly more complex, so we left this for future work.

For patients with multiple seizures available, we obtained the FC matrix for each seizure and then computed the average empirical FC as the mean over all seizures. Figure 2 displays the empirical functional connectivity for the three seizures of Patient 2 together with the SEEG signals selected for each parcel and their amplitude envelope. The amplitude envelope captures the increase in amplitude at seizure onset in the channels with seizure propagation, and the pattern of seizure propagation observed in the empirical SEEG data can be seen in the FC matrix. In Figure 2, the first two seizures display a similar propagation pattern, whereas the third seizure exhibits some differences. Averaging the FC matrices across seizures helps smooth out these variations, capturing the common features of seizure propagation, as shown in the mean FC matrix in Figure 6.

### 2.3 Whole-brain model personalization

#### 2.3.1 NMM description

The personalization pipeline uses one NMM in each node of the network to reproduce the whole-brain dynamics. We have implemented two types of NMM, one for nodes identified by the clinician as belonging to the EZ and another for PZ and NIZ nodes.

In the EZ nodes, located in the parcels indicated by the clinician, we used seizure-generating nodes based on the NMM presented in Lopez-Sola et al. [29], which includes ion dynamics to mimic the transition from the pre-ictal phase to the fast onset phase, and then to the clonic phase. An external input of 110 Hz arriving at the main pyramidal population at 15 seconds forces the model to initiate the seizure at that time point (in contrast to the original work, where the seizures were non-deterministic and depended on the stochastic fluctuations of an external input). The membrane potential, spectral profile, and description of the NMM used in EZ nodes are shown in Figure 3, and the model parameters are summarized in Table A1.

**Figure 3:**
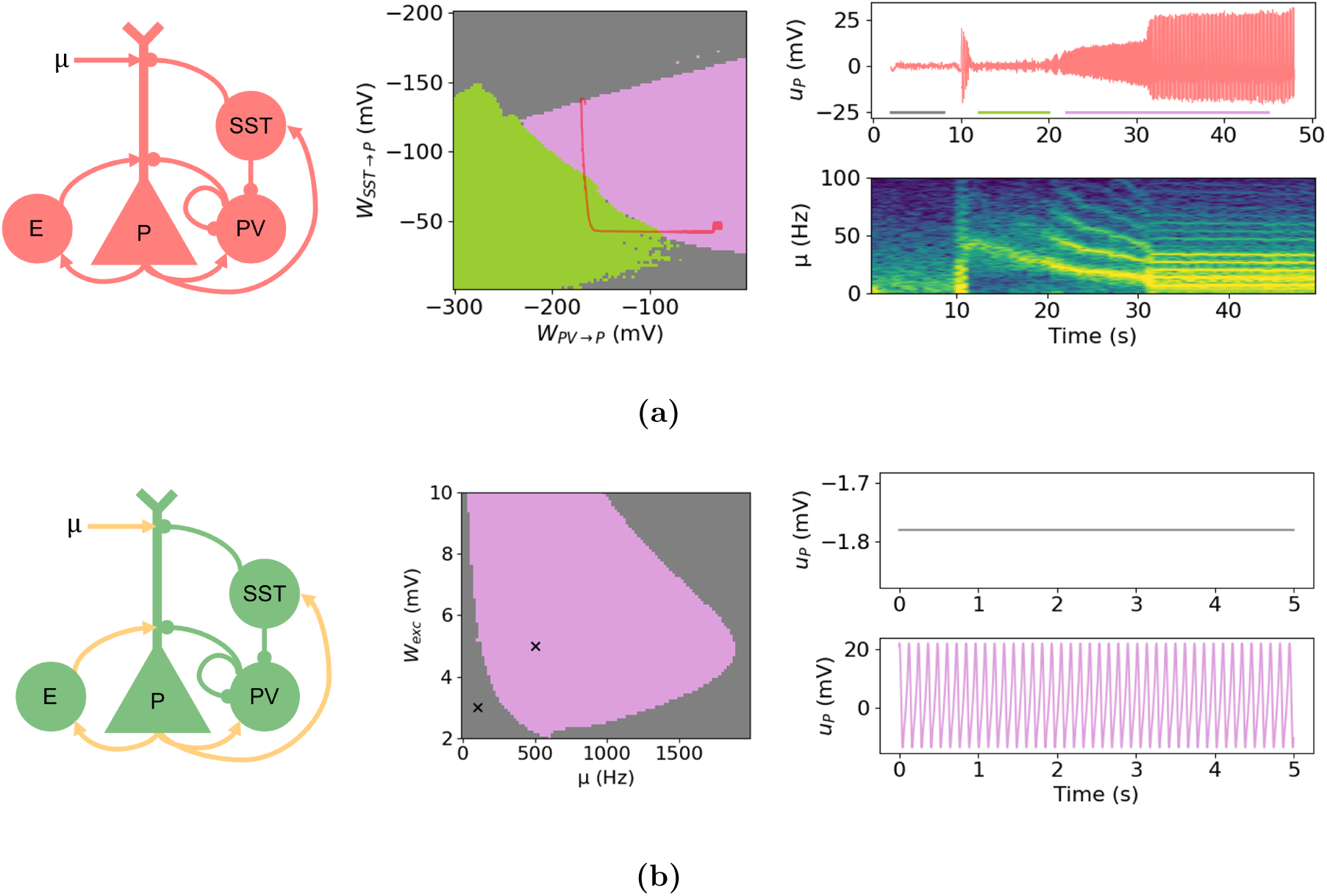
(a) *Left:* EZ NMM architecture (based on [29]). The model includes chloride dynamics in the SST*→*P and PV*→*P synapses to reproduce realistic transitions from the interictal to ictal state. *Middle:* Two-parameter bifurcation diagram showing the different regimes obtained by varying the parameters affected by the chloride dynamics (*W_SST →P_* and *W_P V →P_*). Three main regimes are found: low-voltage noisy dynamics, representing interictal-like dynamics (grey), low-voltage fast activity, representing fast onset dynamics (green), and high-amplitude slow activity, representing ictal-like dynamics (magenta). *Right:* pyramidal population membrane potential in the transition to seizure dynamics and normalized spectrogram. (b) *Left:* PZ/NIZ NMM architecture (based on [20]). The synapses affected by the *W*_exc_ parameter are depicted in yellow. *Middle:* Two-parameter bifurcation diagram obtained by varying the excitatory synaptic gain *W*_exc_ and the external input *µ*. Colors represent the same dynamics as in (a). *Right:* pyramidal population membrane potential for the background regime (top) and the seizure dynamics regime (bottom). Abbreviations: P = pyramidal cells; E = excitatory interneurons; PV = parvalbumin-positive interneurons; SST = somatostatin-positive interneurons; µ= external input to the pyramidal population.

In the PZ and NIZ nodes, a Wendling-class model is used [20]. The specific parameters of the model are taken from [12], as these parameters produce clear transitions from non-seizure to seizure dynamics, with large changes in amplitude when the saddle-node bifurcation is crossed. As will become apparent in the following sections, these amplitude changes are crucial for the current pipeline. The model parameters are summarized in Table A1, and Figure 3 displays the bifurcation diagram of the PZ/NIZ NMM for different values of the excitatory synaptic gain *W*_exc_ and external input *µ*.

When the PZ and NIZ nodes, initially displaying low-voltage non-oscillatory dynamics, receive sufficient input, and depending on the value of the *W*_exc_ parameter, they may cross a saddle-node bifurcation and generate high-amplitude periodic spikes (Figure 3). This transition simulates the propagation of a seizure in PZ and NIZ nodes. To prevent isolated nodes that do not belong to the EZ from seizing spontaneously, we set their mean external input *µ* to 15 Hz (see Table A1). This value lies just below the threshold required to reach the bifurcation point, even for the maximum excitability tested (*W*_exc_ = 10; see Section 2.3.4). As a result, PZ and NIZ nodes can only exhibit ictal dynamics if they receive sufficient input from other active regions, ensuring that seizure propagation in the model is driven by network interactions.

We used a different model for EZ nodes because it captures realistic seizure transitions, including fast onset dynamics, with a single change in external input. In contrast, the Wendling-class model used for PZ/NIZ would require simultaneous changes in multiple parameters to produce comparable transitions, as shown in [20].

#### 2.3.2 Parameters for personalization Global coupling

The nodes in the network are connected based on the structural connectivity of the patient, which is obtained from T1-weighted MRI and diffusion MRI (dMRI) data (see Section 2.2.3). As is common in whole-brain modeling pipelines [7, 17, 52], we have introduced a global coupling gain *G* that multiplies the connectivity matrix and scales the connectivity between nodes. This parameter is personalized for each patient by the optimization algorithm. The range of values used for the *G* parameter is [1, 300], to avoid full disconnection (*G* = 0) and saturation of the neural masses (for *G >* 300 most of the nodes in the network display non-oscillatory activity).

##### Node excitability

We have also personalized for each patient the excitability of the nodes in the network. Although the precise mechanisms of seizure propagation in the brain remain unclear, it is well established that increased excitability of cortical tissue in specific areas is, at least partially, responsible for the initiation and propagation of seizures in the brain [41, 53]. Increased excitability in certain brain areas can result in seizure propagation even when there is weak structural connectivity with epileptogenic nodes. This means that seizure propagation cannot be fully captured by fitting only the global coupling gain *G*, and models personalized using only this parameter resulted in very poor fits.

To implement this physiological property in our models, we have used the excitatory synaptic gain of the neural masses *W*_exc_ as a proxy of the excitability of each brain area (see Table A1 for a complete description of the NMM parameters). For a given node, the personalized value of *W*_exc_ affects all the excitatory synapses targeting the neuronal populations belonging to that node (connections in yellow in Figure 3b), including the excitatory synapses from the external input and from pyramidal populations belonging to other nodes (long-range connections). For nodes with high *W*_exc_ values, the system is closer to the seizure dynamics state. These nodes can enter the region of seizure-like activity even with low values of external input, as shown by the bifurcation diagram in Figure 3b.

The range of values used for *W*_exc_ is [3, 10]. The minimum value, 3 mV, is considered a *healthy* excitability value, as the closest integer to the physiological value used in the Jansen-Rit model [54] (3.25 mV). We chose the maximum excitatory synaptic gain to be 10 mV based on the fact that increasing the excitability of the nodes beyond 10 mV did not substantially increase the probability of the nodes entering seizure dynamics (see Figure 3b).

We personalized *W*_exc_ only for the nodes belonging to regions sampled by SEEG electrodes. For the remaining nodes, we fitted a single parameter, *W*\*_exc_, that represents the excitatory synaptic gain in the nodes where SEEG data was not available. We followed this approach because the model personalization relies on matching the simulated functional connectivity (FC) to the empirical FC derived from SEEG recordings. Empirical FC data is available only for regions in which SEEG electrode contacts are present. Consequently, for nodes outside these regions, fitting the *W*_exc_ parameter has minimal impact on the synthetic FC and introduces degeneracy in the optimization process (risk of overfitting). We can capture the fact that some patients exhibit higher cortical excitability in regions not sampled by SEEG by fitting the *W*\*_exc_ parameter.

#### 2.3.3 Simulated SEEG and FC

As in our previous studies [29–31, 55, 56], we have added a laminar physical model to the NMM formalism to generate realistic physiological measurements, in this case SEEG. In particular, we have used the modeling work presented in [31], which combines the laminar NMM framework and a geometric model of synaptic currents, to simulate the current source density (CSD) inside a cortical patch and derive the induced voltage recorded by a pair of adjacent SEEG contacts.

##### CSD computation from NMM outputs

In brief, we used the NMM’s output to calculate the average post-synaptic current, *I_s_*(*t*), associated with each of the synapses from other populations into pyramidal cells. We have considered these neurons as the main contributors to the measured voltage, given the anatomical characteristics and organization of pyramidal cells [57, 58]. Each of these currents is proportional to the average membrane potential perturbation at each neuron, *u_s_*, induced by the associated pre-synaptic population, *I* = *η · u_s_*, with *η* a gain proportionality factor of units A/V. We have used a gain factor of *η* = 10^−12^ A/mV to generate SEEG signals of the same magnitude as the empirical ones.

The current source density depth profile over time *CSD*(*z, t*) can be generated from the average post-synaptic currents computed above using the cortical patch model and the distribution of synapses in cortical layers presented in [31]. The CSD corresponding to the activity of the EZ NMM in one parcel is shown in Figure B1.

##### Generation of SEEG from CSD

Next, we have used the current source density profile *CSD*(*z, t*) to calculate the voltage measured by a pair of SEEG contacts *V_SEEG_*(*z_el_*) as described in [31]. As demonstrated in that work, and as shown in Figure B1, the magnitude of the synthetic SEEG signals depends on the position of the electrode contacts. In [31], SEEG contacts were modeled as 0.8 mm diameter and 2 mm length cylinders separated by an insulating material with 1.5 mm of length. Given that the precision about the electrode depth from the imaging data is very low compared to the sensitivity of the SEEG measurements as a function of the electrode depth, we have assumed the same electrode depth for all nodes. We have chosen a value of *z_el_* = *−*1 mm, which corresponds to both electrode contacts equally inside the grey matter layer.

It is worth mentioning that other studies have used different paradigms to simulate SEEG signals or local field potentials (LFPs), notably NMM-derived dipole activity or the sum of excitatory and inhibitory PSPs at the level of the main pyramidal population [59, 60]. Although the current framework is thought to better represent the connection between the neural mass model formalism and electrophysiological measurements, the choice of the physical model (or the depth of the SEEG electrode in our physical modeling) is not expected to substantially change the results of the personalization pipeline. This is because the loss function for personalization is based on changes in signal amplitude, and we expect abrupt changes in the synaptic activity of a given NMM when the input received is sufficiently large due to the presence of a bifurcation. The translation from NMM synaptic activity to physiological measurements is expected to preserve these changes in amplitude, independently of the physical modeling used to generate such measurements.

##### Noise and filtering

We added noise to the SEEG signals generated to replicate the background noise measured in empirical data. We found that adding 1/*f*^2^ noise replicates well the power spectral density (PSD) profile observed in real data, and adjusted the magnitude of the noise so that the PSD scales of synthetic and empirical data were reasonably similar (see Figure B2). Finally, we filtered synthetic SEEG signals between 0.5 Hz and half of the sampling frequency (1000 Hz for simulated data, 512 or 1024 Hz for empirical data) to reproduce the usual hardware filtering of empirical SEEG data.

##### Computation of synthetic FC

The calculation of the FC between synthetic SEEG signals was performed in the same way as for empirical SEEG data. We used the maximum cross-correlation between the amplitude envelope of synthetic SEEG signals in a window defined by the start of the clonic phase. The detection of the clonic phase in synthetic signals is based on the firing rate of the excitatory population (E) in each NMM, which displays high-amplitude spikes that saturate (they reach the maximum firing rate of 5 Hz) when the system enters seizure-like dynamics. The presence of such spikes was selected as the beginning of the clonic phase in the NMM.

#### 2.3.4 Loss function for personalization

We defined the loss function to be minimized by the optimization algorithm as:

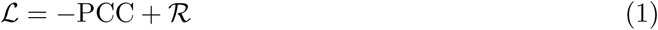

PCC represents Pearson’s correlation coefficient between the empirical and synthetic FC matrices, and *R* is a regularization term that captures the deviation of the parameter *W*_exc_ from healthy values (see Equation 2). Thus, by minimizing *L*, we will find parameter solutions that maximize the similarity between the seizure propagation pattern in empirical and simulated data while minimizing the excitability in the network nodes.

##### L1 regularization

To address the possible degeneracy when fitting the excitatory gain parameters *W*_exc_, particularly in nodes that have little influence on the SEEG-derived FC, we introduced a regularization term in the loss function (Equation 1). Assuming that most brain areas are healthy, we can consider them to be close to a baseline excitability value *W*_exc, h_, while only a few regions exhibit pathological hyperexcitability. To reflect this sparsity, we applied L1 regularization to penalize deviations from the healthy baseline,

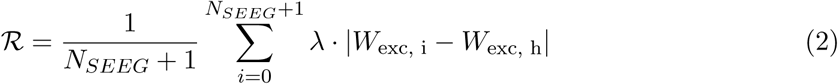

In Equation 2, *W*_exc, h_ = 3 represents the healthy baseline excitability (see Section 2.3.2), *N_SEEG_* is the number of nodes in regions sampled by SEEG electrodes, and *λ* is a regularization parameter. Since the number of SEEG-sampled nodes varies across patients, we normalize the penalty by *N_SEEG_* +1, where the additional 1 accounts for the extra excitability parameter *W*\*_exc_ used for nodes belonging to regions not sampled by SEEG. This normalization ensures that the strength of the regularization remains comparable across patients regardless of the number of fitted parameters. The value of *λ* was selected to promote sparsity while preserving fit quality, as measured by the PCC between empirical and synthetic FC (see Figure C1).

##### Model validity

We restricted the search to models that are in the desired dynamical regime and do not reproduce unrealistic physiological activity. In particular, we have considered models as valid if the following criteria were met:

- A seizure is present in all EZ nodes.
- The clonic phase in EZ nodes doesn’t start before 5 seconds. The clonic phase detection was performed as detailed in Section 2.3.3. This ensures that the seizure transition from pre-ictal to fast onset to clonic is preserved.
- Seizures are not detected in PZ or NIZ nodes before the clonic phase starts in the EZ nodes. This ensures that PZ or NIZ nodes do not display seizure activity if it’s not due to seizure propagation.
- The difference between the minimum value of synthetic FC and the minimum empirical FC is not above 0.2. This avoids fittings where the empirical and synthetic FC have a high PCC but their scales are very different. In particular, we want to avoid situations where all the synthetic signals are highly correlated (seizure activity in all nodes), but small differences in the signals result in patterns similar to those of the empirical FC matrix.

#### 2.3.5 Optimization algorithm

The number of parameters to fit is in the order of 10–30, depending on the number of VEP parcels sampled by SEEG electrodes. We have used the differential evolution (DE) algorithm [61] to solve the optimization problem, in particular, the algorithm implementation provided by the SciPy Python library using the *best1bin* strategy [62].

We selected the values for the mutation rate, recombination rate, population size, and maximum number of generations used in the differential evolution algorithm to balance fitting performance and computational efficiency, as detailed in Appendix C.2. To reduce the search space and improve convergence, we restricted the parameter search to integer values.

The initialization of the population is a critical step to ensure effective exploration of the parameter space and improve the likelihood of finding the global minimum. For the global coupling parameter *G*, values for each individual in the initial population were drawn from a uniform distribution spanning the allowed range (0 to 300). For the excitability parameters *W*_exc_, we adopted an initialization strategy aligned with the L1 regularization used in the loss function, which assumes a Laplacian prior centered at the healthy value *W*_exc, h_ = 3 (see Equation 2). Since 3 is also the lower bound for *W*_exc_, we used a “half-Laplacian” distribution that peaks at this minimum and decays for higher values. Each initial *W*_exc_ value (including the global non-SEEG parameter *W*\*_exc_) is generated as

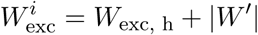

where *W*’ is drawn from a Laplacian distribution centered at 0 with a scale parameter of 1. This approach biases the initialization toward healthy excitability while allowing for exploration of pathological values.

Each whole-brain simulation was run for a total duration of 50 seconds using a sampling frequency of 1000 Hz with a Runge-Kutta 4 solver.

### 2.4 Evaluation of personalization results

To evaluate the success of the personalization for each patient, we computed the Pearson correlation coefficient (PCC) between the empirical and simulated functional connectivity (FC) matrices, using it as a measure of goodness-of-fit. As a baseline, we also computed the PCC between the empirical FC and the structural connectivity to confirm that the model could capture functional dynamics beyond what is directly encoded in the connectome.

To further assess the physiological plausibility of the personalized models, we compared the presence of seizures in the empirical and simulated SEEG signals. For simulated data, seizure detection was performed using the method described in Section 2.3.3. For empirical data, we implemented an energy ratio–based detection approach. For each bipolar contact, we bandpass filtered the signal (1–50 Hz) and computed the energy ratio between the seizure window (50 seconds post-onset, as marked by the clinician) and the preceding baseline. Energy values were normalized by the maximum ratio across contacts, and a fixed threshold of 10% was used to classify each region as presenting seizures or not.

To quantitatively compare the empirical and simulated seizure patterns, we computed the Jaccard index between the sets of regions with detected seizures, as a measure of spatial overlap of active nodes in real and simulated data. In addition, we constructed confusion matrices for each patient and derived standard performance metrics, including sensitivity, specificity, and F1-score.

### 2.5 Simulation of interventions

#### 2.5.1 Node removal

To evaluate the potential of the model to predict the outcome of surgery-like interventions, we studied the effect of removing individual nodes from the network. Although implemented as node removal in the model, this manipulation can be interpreted as mimicking either surgical resection or functional disruption through neuromodulation, such as tDCS. These interventions differ in practice, but both aim to reduce seizure initiation and propagation by targeting key regions in the network.

To remove a given node, we removed from the whole-brain model parameters all the neurons and synapses associated with this node. We then simulated the whole-brain model with the new NMM architecture. We only removed nodes that belonged to parcels sampled by SEEG and that exhibited seizure propagation in the original model, as these were the ones contributing to seizure spread and thus whose removal could reduce it. To understand which nodes would lead to better surgical outcomes, we compared the number of nodes exhibiting seizures in the model before and after node removal. Seizure detection in the model was performed as detailed in Section 2.3.3.

#### 2.5.2 Pharmacological interventions

We simulated three types of pharmacological interventions in the model to evaluate patient-specific responses to different treatments: benzodiazepines, stiripentol, and felbamate. Each drug was modeled by modifying a specific parameter of the neural mass model, based on known mechanisms of action:

- **Benzodiazepines** enhance GABA_A_-mediated synaptic amplitude (GABA_A_ agonists). To simulate their effect, we increased the synaptic gain of inhibitory synapses (*W*_inh_), following previous work [22].
- **Stiripentol** prolongs the opening duration of chloride channels on GABA_A_ receptors (GABA_A_ modulator). We modeled this by increasing the inhibitory synaptic time constant (*τ*_inh_), similarly to previous studies [22].
- **Felbamate** is an NMDA receptor antagonist. Its effect was modeled by reducing the synaptic gain of excitatory synapses (*W*_exc_), acting as a counterbalance to increased local excitatory connectivity in certain brain regions.

We modeled drug effects using plasma concentrations expressed in mg/L. For each drug, we defined three concentration levels (low, medium, and high) corresponding to 1%, 5%, and 10% changes in the respective model parameter. We selected these percentages based on preliminary simulations, where a 10% change often led to complete seizure suppression in many patients. The associated concentrations reflect clinically relevant therapeutic ranges: (1) benzodiazepines (clonazepam): 0.02, 0.05, and 0.07 mg/L; (2) stiripentol: 4, 13, and 22 mg/L; and (3) felbamate: 30, 45, and 60 mg/L [63, 64].

In all cases, parameter changes were applied uniformly across all nodes as a percentage of the original values in each region. While *W*_inh_ and *τ*_inh_ were constant across parcels, *W*_exc_ varied spatially as part of model personalization.

## 3 Results

### 3.1 Personalized models replicate the pattern of seizure propagation

We first analyzed the success of the personalization for each patient by computing the PCC between the empirical and synthetic FC matrices of each patient, as a measure of goodness-of-fit. We compared such PCC with the PCC between the patients’ empirical FC and structural connectivity, to confirm that the model could reproduce functional dynamics beyond the information provided by the connectome. Figure 4 shows the personalization results for the six patients included in this study. In all cases, the PCC between the empirical and synthetic FC matrices is higher than the PCC between the empirical FC and the structural connectivity, with a mean correlation of 0.55. We confirmed that the personalized models showed significantly higher correspondence with empirical FC by statistically comparing the model-derived PCC values to those obtained using structural connectivity alone (p=0.031, Wilcoxon signed-rank test).

**Figure 4:**
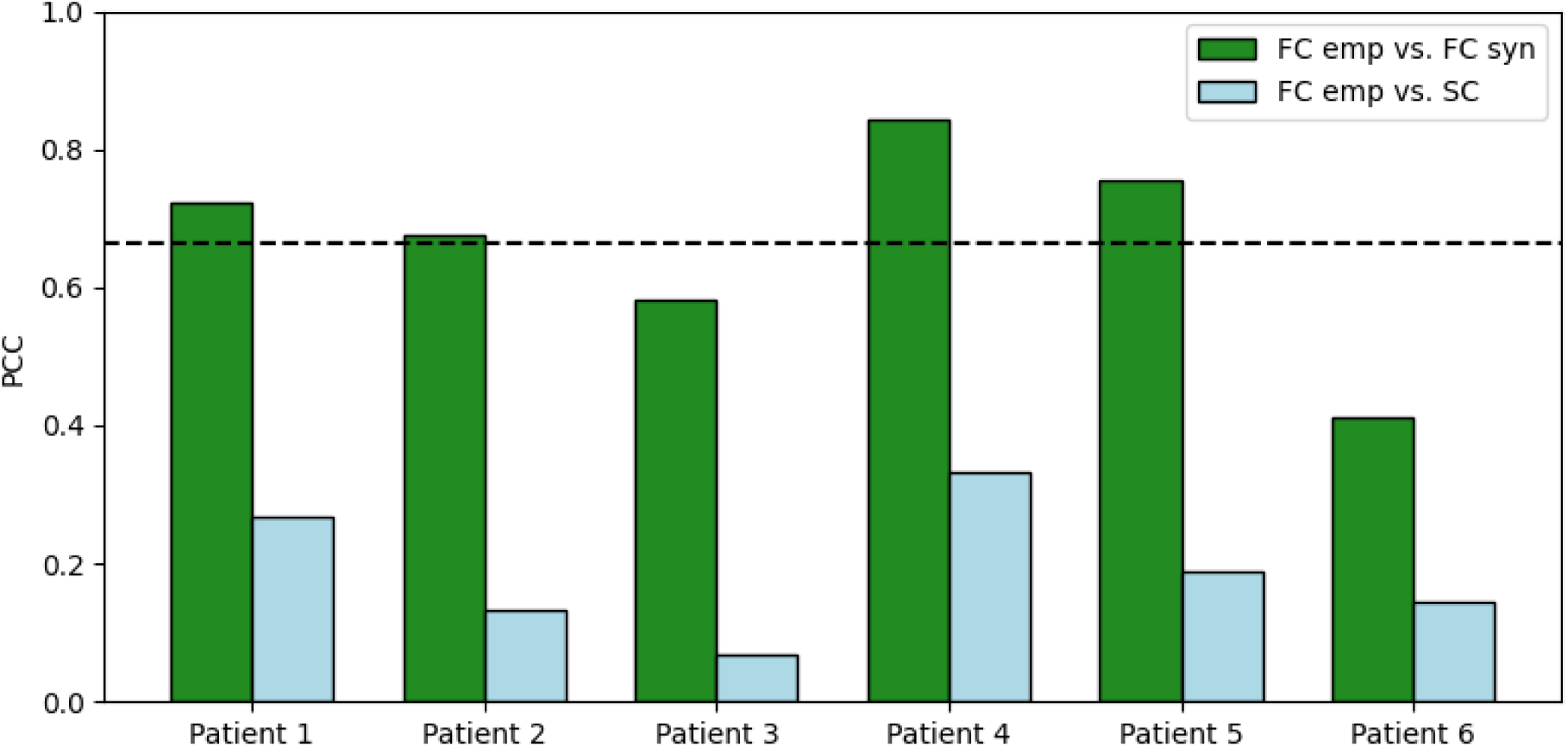
Personalization results: comparison of PCC between the empirical and synthetic FC (target for model fitting) and the PCC between the empirical FC and the structural connectivity (as a quality check). The dashed line indicates the average PCC between empirical and synthetic FC.

We also assessed the robustness of the personalization by comparing the presence of seizures in the empirical SEEG data to the presence of seizures in the synthetic SEEG signals. The personalized whole-brain models match the pattern of seizure propagation, as illustrated by the comparison of empirical and simulated SEEG signals (Figures 5 and D2) and of the empirical and synthetic FC matrices (Figure 6). The relatively stationary tonic firing observed in PZ and NIZ nodes reflects the simplified nature of the underlying model, which is designed to capture seizure recruitment and propagation rather than detailed temporal features such as bursting or intermittent dynamics. The complete set of empirical and synthetic SEEG signals, as well as the values of the personalized parameters for each patient, is provided in Appendix D.

**Figure 5:**
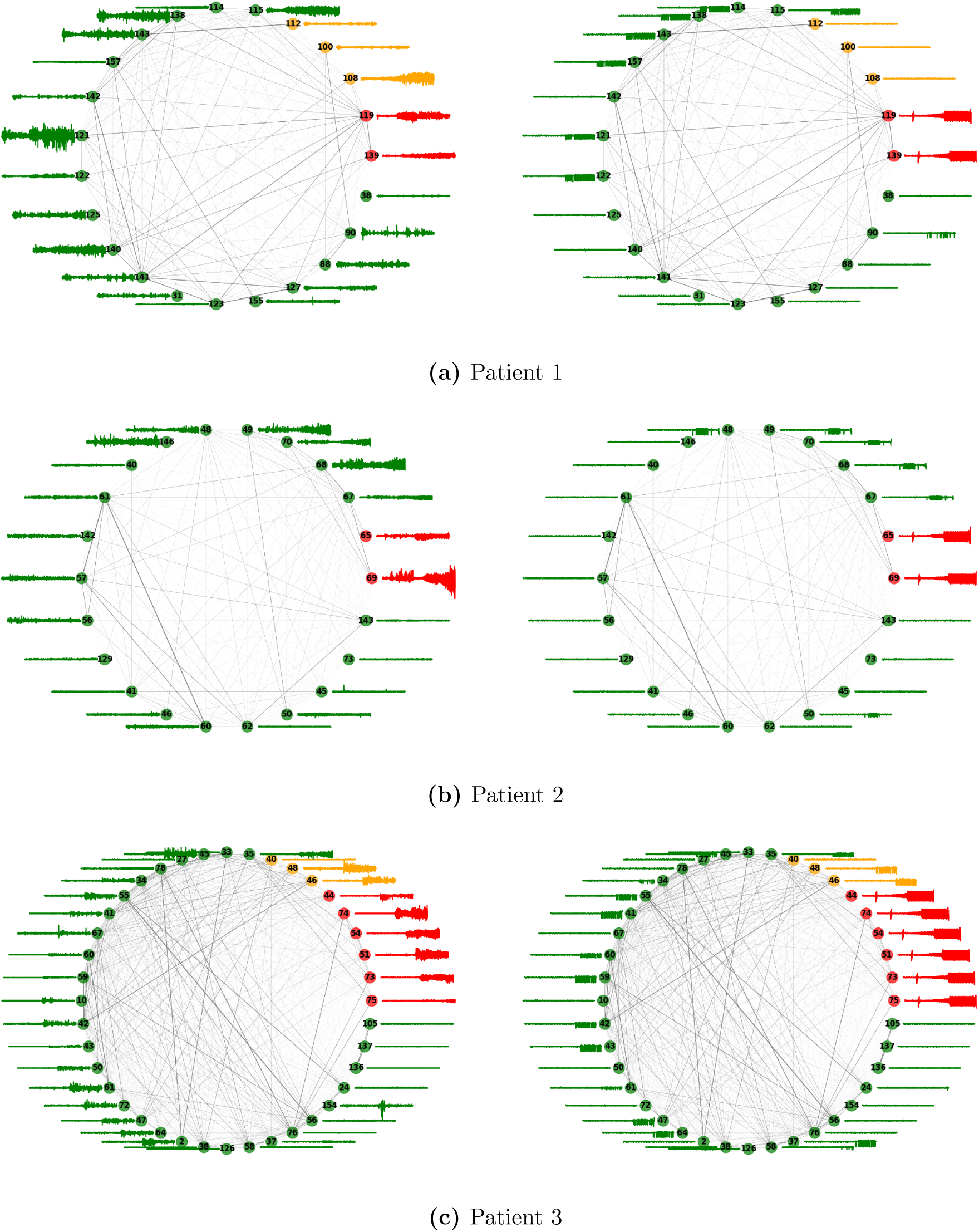
Comparison between empirical (left) and synthetic (right) SEEG signals for three patients of the study. For the empirical data, only one seizure recording is shown. Node numbers represent parcel numbers in the VEP parcellation, and edge width represents the connectivity strength between two nodes. Colors indicate the epileptogenicity of the regions: red = EZ, yellow = PZ, green = NIZ. SEEG signals are shown without axis labels for clarity, the range displayed is from –1000 to 1000 µV. The rest of the patients are shown in Appendix D.

**Figure 6:**
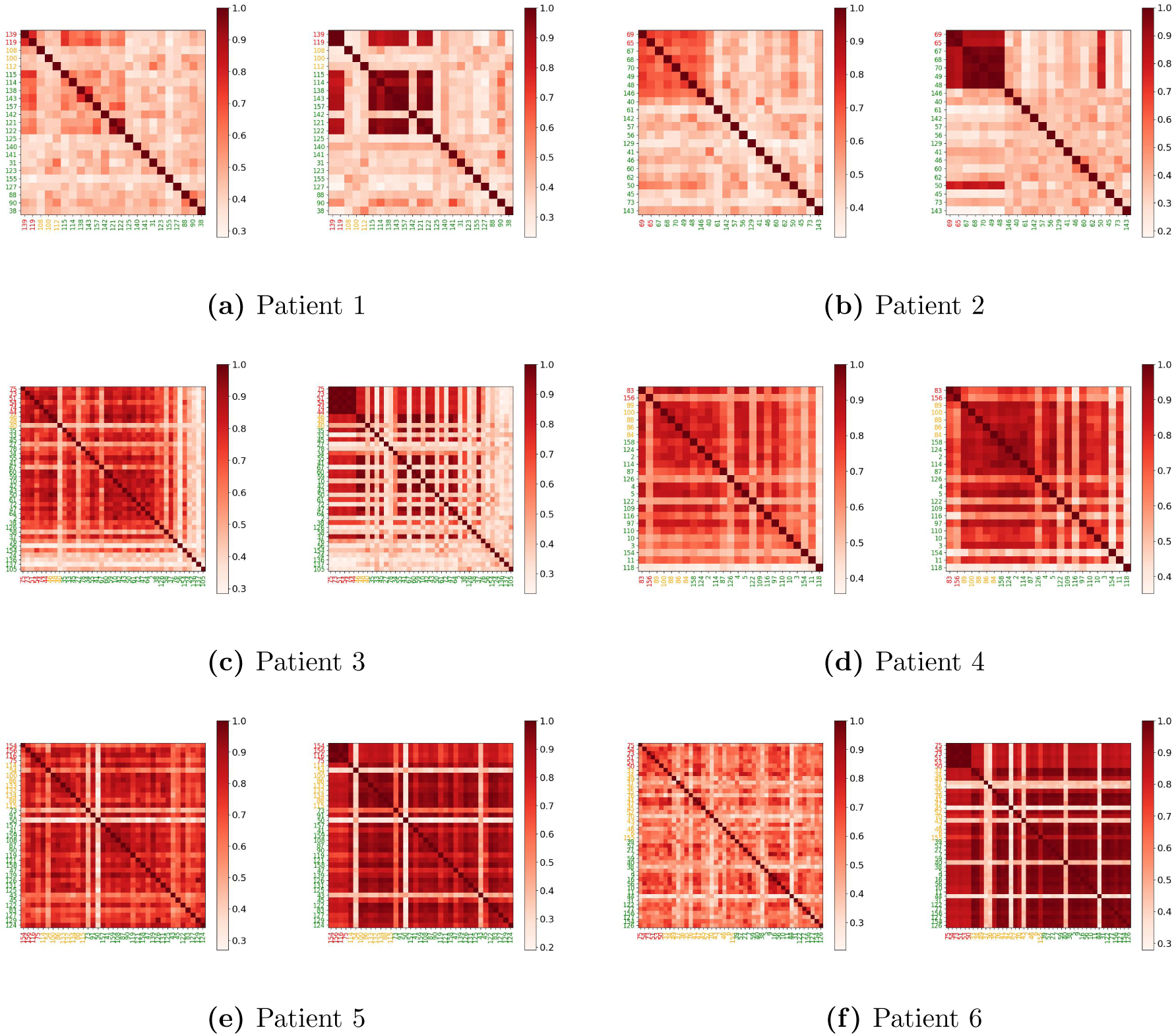
Empirical (average) FC (left) and synthetic FC after model personalization (right) for each of the six patients in the study. Colorbars represent PCC values, and region labels are colored according to clinical classification: red = epileptogenic zone (EZ), orange = propagation zone (PZ), green = non-involved zone (NIZ).

To quantitatively assess how well the model reproduced the spatial distribution of seizure activity, we compared the regions where seizures were detected in the empirical and simulated SEEG data. We computed the Jaccard index between the empirical and simulated sets, as well as standard classification metrics derived from confusion matrices. Across patients, the average Jaccard index was 0.59 (SD = 0.10), indicating substantial spatial overlap. The mean sensitivity was 0.89 (SD = 0.13), with an F1-score of 0.74 (SD = 0.07), showing that the models reliably captured core seizure regions, albeit with some overprediction, as reflected in a lower specificity (mean = 0.65, SD = 0.10). Full detection maps and confusion matrices for each patient are shown in Appendix D.

Interestingly, for some subjects, the whole-brain models predict seizure propagation in areas not explored by the SEEG electrodes. Figure D4 shows the nodes that display seizure in each of the whole-brain model simulations. We can see that for three of the six patients in the study, the model predicts widespread seizure propagation. We could confirm that for those patients the empirical SEEG data displayed seizure-like activity in most of the nodes sampled (see Figures 5 and D2). The integration of scalp EEG seizure data into the personalized models would confirm the validity of such predictions, a possibility we discuss in Section 4.2.

### 3.2 Node removal reveals candidates for interventions

We studied the effect of removing individual nodes from the network and measured the total number of nodes exhibiting seizures after each removal. Figure 7 shows the results of this analysis. In each case, certain nodes appear to be critical for seizure propagation and may be promising targets for interventions such as surgery or tDCS.

**Figure 7:**
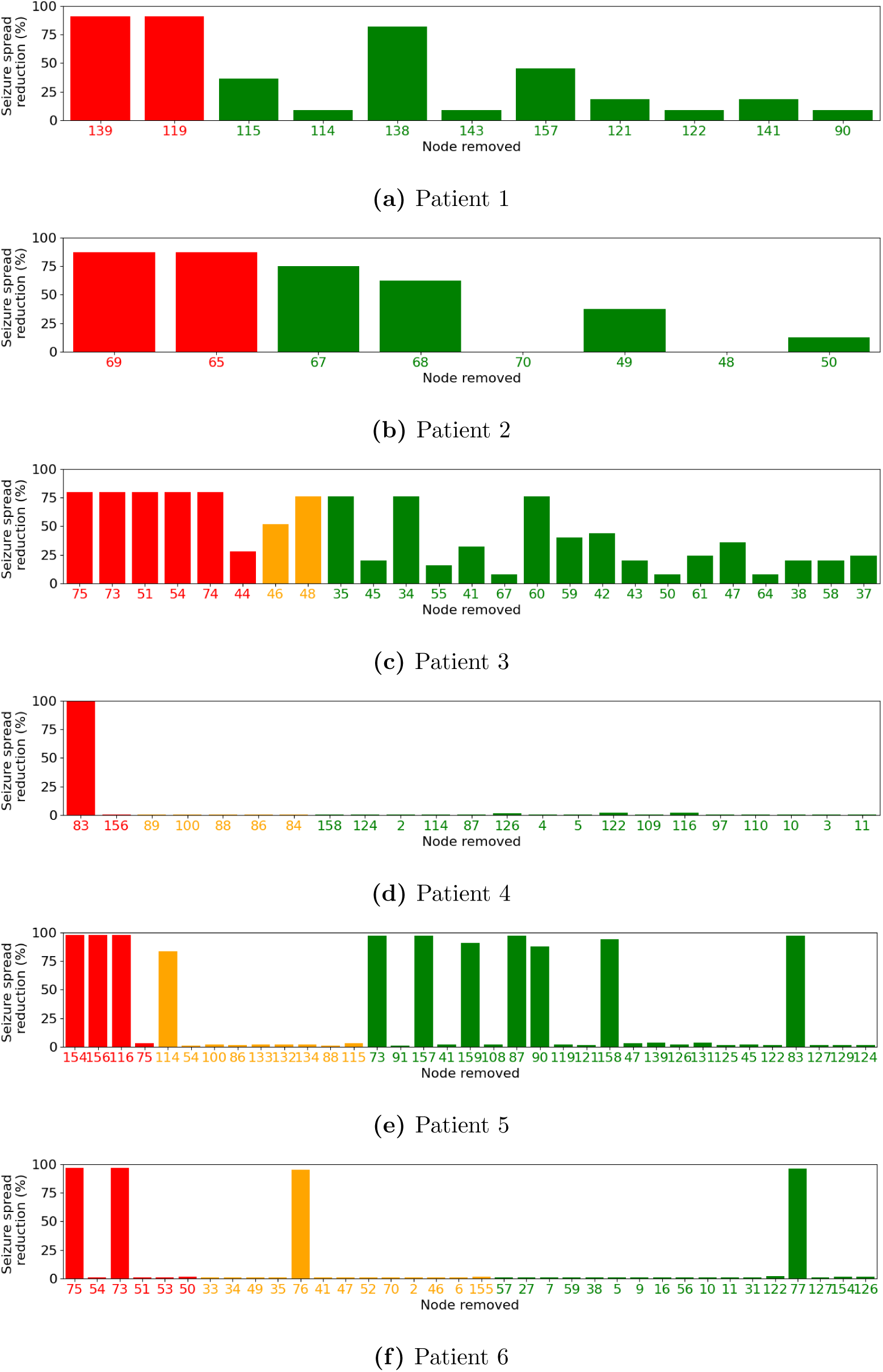
Node removal. Reduction of seizure spread (in %) depending on the node removed from the network. We considered for removal only those nodes sampled by SEEG electrodes displaying seizure activity. Colors indicate the epileptogenicity of the regions: red = EZ, yellow = PZ, green = NIZ.

As expected, removing EZ nodes leads in most cases to a reduction in seizure spread, and in some cases even to complete seizure suppression. However, not all EZ nodes contribute equally to propagation. Some appear to play a more central role, as their removal results in a more pronounced reduction of seizure activity. For example, in Patient 4, removal of EZ node 83 leads to a greater reduction in seizure spread than removal of EZ node 156. Note that in our model, EZ nodes are deterministically driven into seizure by an external input and cannot be suppressed by removing other regions. As a result, competitive interactions between multiple EZs are not captured here.

Interestingly, the model also predicts that removing certain PZ or NIZ nodes can reduce seizure propagation. While the effect is generally smaller compared to removing EZ nodes, in some cases, it is still substantial. For instance, in Patient 2, removal of nodes 67 and 68, which are not part of the EZ, led to a noticeable decrease in seizure spread. These nodes showed seizure activity both in the model and in the empirical SEEG data (see Figure 5), suggesting that they may play a secondary but functionally relevant role in the propagation network.

We also observe variability across patients in the number and type of critical nodes, reflecting differences in individual network organization and seizure dynamics. In some patients, multiple nodes significantly contribute to seizure propagation (e.g., Patient 3), while in others (e.g., Patient 4), seizure spread appears to depend on a single dominant node. This highlights the importance of personalized modeling for identifying optimal intervention targets, which may not always coincide with the clinically defined EZ.

To further explore the relationship between the critical nodes predicted by the model and the EZ regions identified clinically, we examined the anatomical locations of the most impactful nodes for selected patients. In Patient 1, the EZ was located in the right supramarginal gyrus (anterior) and the right planum temporale (T1), and the non-EZ node that led to the greatest seizure reduction was the right insula (gyri longi), a region anatomically adjacent to the supramarginal and posterior temporal areas. In Patient 2, the EZ involved the left occipital pole and left anterior occipital sulcus, and the most critical non-EZ nodes were located in adjacent visual areas, including left O2 and O1, reflecting anatomical and functional continuity within the occipital lobe. In Patient 3, the EZ included mesial temporal lobe structures such as the left amygdala and hippocampus. The top non-EZ critical nodes were in the T2-posterior and T3-posterior regions, located in the posterior inferior temporal cortex. These regions are anatomically connected to the mesial temporal lobe and represent typical pathways of seizure propagation in temporal lobe epilepsy, supporting the relevance of the targets predicted by the model. These examples suggest that the model tends to identify critical regions that are closely related to the EZ regions defined clinically in terms of anatomical proximity and known propagation routes.

It is important to note that while some model predictions suggest almost complete seizure suppression following the removal of specific nodes (including PZ and NIZ nodes in Patients 4, 5, and 6), we do not interpret these results literally. It is, in practice, unlikely that removing a single node, particularly one outside the EZ, would completely eliminate seizure initiation. This outcome arises in the model because seizure activity is driven by interactions across the network: disrupting key propagation pathways can, in some cases, prevent other nodes from reaching the threshold for seizure propagation. Therefore, while these predictions of seizure suppression should be interpreted cautiously, the analysis still provides valuable insights into the relative importance of different nodes in the propagation network and can help identify secondary targets whose modulation may reduce seizure spread.

### 3.3 Simulation of pharmacological interventions

We studied the effect of different pharmacological interventions on network seizures in the models. Figure 8 shows the impact of the three treatments tested.

**Figure 8:**
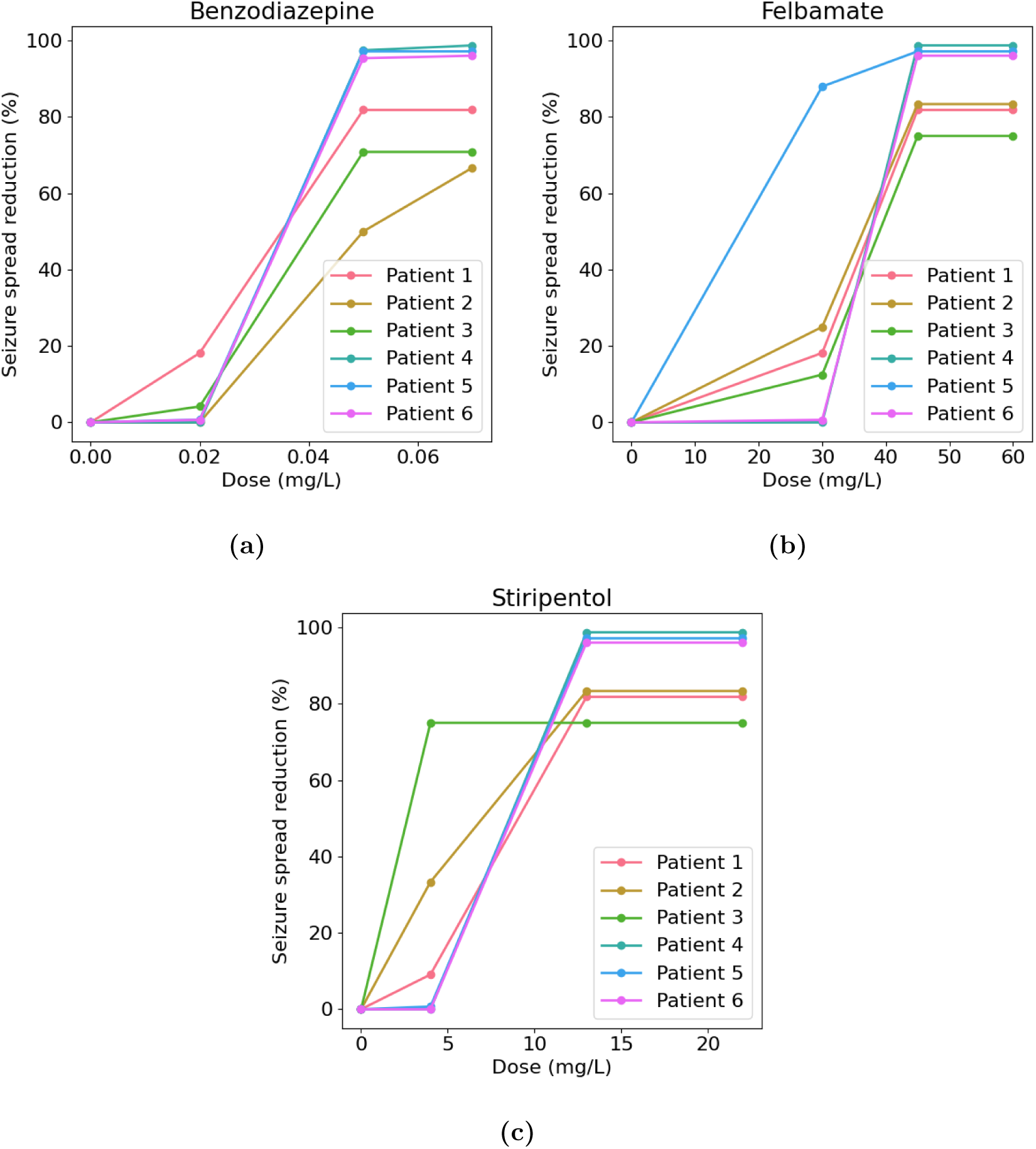
Pharmacological intervention. Reduction of seizure spread (in %) for different doses of different pharmacological agents: (a) benzodiazepines, (b) stiripentol, and (c) felbamate. Three concentration levels (low, medium, and high) were considered, corresponding to 1%, 5%, and 10% changes in the respective model parameter. The selected concentrations reflect clinically relevant therapeutic ranges [63, 64].

The personalized whole-brain models predict varying patient responses to pharmacological interventions, highlighting the importance of individual variability in treatment efficacy. While most patients show a dose-dependent reduction in seizure spread, the degree and nature of the response differ across drugs and individuals. For instance, Patient 3 displays a strong response to stiripentol (a GABAergic modulator), suggesting that enhancing inhibitory tone is particularly effective in this case. In contrast, Patient 5 responds more markedly to felbamate, an NMDA receptor antagonist, indicating a possibly dominant role of excitatory drive in seizure propagation for this patient.

Interestingly, for some individuals (e.g., Patient 1), the models suggest that multiple drug classes could lead to seizure suppression, albeit to different extents. In others (e.g., Patient 6), the response is robust across all drugs at higher doses, but limited at lower concentrations, possibly reflecting a higher excitability threshold for pharmacological modulation.

## 4 Discussion

Our results demonstrate that it is possible to successfully personalize whole-brain computational models using patient-specific structural and functional data, resulting in individualized simulations that replicate observed seizure propagation patterns. Importantly, these personalized models yield distinct responses to virtual interventions (both surgical, pharmacological, or resulting from local brain stimulation [65]), highlighting their potential for guiding patient-specific treatment strategies. While these findings are promising, the translation to clinical application will require thorough validation of this pipeline. A natural next step would be a retrospective analysis of surgical cases, comparing model predictions regarding critical propagation nodes with actual surgical resections and outcomes. Such validation could help assess the predictive power of the models and their potential role in pre-surgical planning or intervention design [18, 43].

The dynamics of seizure initiation and propagation in our models can be interpreted using bifurcation theory: each node operates near a threshold where changes in excitability or input can trigger seizure activity. Excitability, modulated by *W*_exc_, interacts with incoming network input to determine a node’s state. During personalization, *W*_exc_ is tuned so that the network reproduces the empirically observed propagation pattern. Crucially, a node’s seizure state depends not only on its local properties but also on inputs from other regions. This explains why removing a single node can prevent downstream regions from seizing. While predictions of seizure suppression from PZ or NIZ nodes may seem overly optimistic compared to clinical observations, they reveal hidden hubs in the propagation network and suggest indirect targets for intervention. Further validation with additional data (such as scalp EEG) and clinical outcomes will be key to confirming these predictions.

Unlike more abstract models such as the Epileptor [6], which are tailored to capture the dynamics of seizure initiation and termination but lack a clear link to the underlying physiology, our neural mass modeling approach enables the simulation of diverse interventions within a unified framework. This is due to the model’s explicit physiological parameters, such as excitatory synaptic gain and receptor-specific conductances, which can be directly manipulated to mimic the effects of pharmacological treatments or neuromodulation. The results presented here illustrate this capability: we simulated both node removal (as a proxy for surgical intervention or stimulation) and drug modulation by altering specific biophysical parameters, and observed how these interventions affected seizure dynamics across patients. The fact that distinct patterns of response emerged across interventions and individuals underscores the versatility of our approach and supports its use as a tool for exploring treatment strategies beyond seizure localization.

Although all patients included in this study are clinically classified as drug-resistant, our models still predict responses to certain pharmacological interventions. This is not necessarily inconsistent: drug resistance is typically defined after failure of two or more appropriately chosen and dosed medications [1], but does not imply that all potential drug combinations or concentrations have been tested. While we did not have access to detailed records of the medications each patient received, limiting direct comparison with clinical outcomes, our framework provides a mechanistic way to explore alternative pharmacological or combined strategies that might be effective but were not considered or optimized in clinical practice. As such, it may help refine drug selection or dosing in a personalized manner, potentially discovering treatment options that would otherwise be overlooked.

### 4.1 Limitations

A key limitation of the current modeling pipeline is that the fitting process relies exclusively on SEEG data, which samples only a limited portion of the brain. Although the personalized models can make predictions about seizure propagation beyond the regions directly observed with SEEG, the accuracy of these predictions is constrained by the sparse and potentially biased spatial sampling. To address this limitation and improve model personalization, future work will integrate scalp EEG data, which offers broader coverage of cortical activity [60]. The combination of SEEG and EEG will enable us to better constrain the model dynamics in both deeply and superficially located brain regions, ultimately enhancing the robustness and generalizability of the personalized models.

Another limitation of the current pipeline is that model personalization relies on the average of empirical functional connectivity (FC) matrices across seizures. While this approach is effective in patients with consistent seizure propagation patterns, it may fail to capture the full variability present in cases where seizures follow distinct propagation routes. In such scenarios, the averaged FC matrix may obscure critical features of individual seizure dynamics, potentially leading to a misrepresentation of the true underlying propagation network. As a result, the personalized model might reproduce a generalized propagation pattern that includes all regions involved across seizures, but not necessarily the specific dynamics of any single event. Although this can still be valuable for identifying common targets for intervention, it may limit the precision of predictions in patients with high seizure heterogeneity.

It is worth mentioning that most epilepsy patients, but not all, display seizure onset patterns that include a clonic phase characterized by an amplitude increase [50]. In our case, all patients studied presented this characteristic. Nevertheless, it is possible that the current pipeline could still perform reasonably well in cases without such a pronounced amplitude increase at the onset of the seizure. For instance, many cases include pre-ictal discharges [50], and those can be captured by the amplitude envelope method used to compute the FC. In such cases, the pattern of correlation shown in the FC matrix could correspond to the propagation of the pre-ictal spikes, which are thought to be generated by mechanisms related to epileptogenesis, distinct from those generating inter-ictal discharges [66]. In the rare cases where both the amplitude increase of the clonic phase and the pre-ictal discharges are absent, the personalization algorithm presented here might fail to capture the propagation of seizure dynamics. An alternative approach in such cases could involve identifying fast-onset activity patterns, which have been proposed as reliable markers of seizure onset [67].

Another limitation of the current approach is the assumption that drug effects are applied uniformly across all brain regions. While this simplifies model implementation, it may overlook the spatial heterogeneity arising from factors such as regional differences in receptor density or pharmacokinetics. Although the receptors targeted by the drugs modeled in this work (GABA and NMDA receptors) tend to have relatively widespread distributions, future work could explore whether incorporating spatially varying drug effects improves prediction accuracy or alters intervention outcomes.

### 4.2 Future work

In future work, we aim to extend the model validation beyond ictal activity by evaluating how the personalized models reproduce the interictal state. While seizure data provides critical information for identifying seizure propagation pathways, interictal SEEG recordings are more abundant and can offer valuable insights into the underlying network dynamics in the absence of seizures. By comparing model-generated activity with interictal SEEG data, particularly in terms of spatial distribution of interictal discharges, we can further assess the physiological plausibility and stability of the personalized models. This will also help determine whether seizure-prone regions exhibit aberrant dynamics even in the interictal period, potentially enhancing the models’ predictive power. As discussed, we also plan to incorporate scalp EEG data into the pipeline to improve spatial coverage and further constrain model dynamics.

Another promising direction for future work involves the development of more sophisticated algorithms for therapeutic intervention planning. In this study, we explored the effects of individual node removal to simulate surgical resection and understand the influence of specific regions on seizure propagation. We focused on single-node interventions due to the computational cost of simulating all possible combinations, but expanding the model to include multi-site resection or stimulation could reveal synergistic effects and support the design of more effective and minimally invasive strategies.

Furthermore, the current approach does not capture the full potential of neuromodulation strategies such as tDCS or DBS, which can dynamically modulate activity without permanently altering brain structure. For instance, DBS may suppress seizures by disrupting pathological synchronization across distributed regions, while tDCS can influence a broader spatial extent through diffuse electric fields. We are currently developing optimization algorithms that can identify optimal stimulation targets and protocols, such as multi-channel tDCS, to better reduce seizure propagation [68, 69]. Incorporating plasticity mechanisms is also essential to capture the long-term reorganization effects of such treatments and better understand their sustained therapeutic impact [70].

## 5 Conclusions

In this study, we showed that personalized whole-brain models could reproduce patient-specific seizure propagation patterns using SEEG, MRI, and dMRI data. These models were able to match the seizure propagation observed in SEEG data, as captured by the functional connectivity between SEEG signals. The personalized models provided maps of excitability that varied across brain regions and predicted propagation of seizures in areas not sampled by SEEG electrodes. We also demonstrated how these models can be used to simulate different types of interventions, including surgery, tDCS, and pharmacological treatments. These findings support the use of personalized whole-brain modeling as a framework for understanding seizure propagation and exploring treatment options in refractory epilepsy.

## Acknowledgements

This work has received funding from the European Research Council (ERC) under the European Union’s Horizon 2020 research and innovation programme (Grant Agreement No. 855109; ERC-SyG 2019) and from FET under the European Union’s Horizon 2020 research and innovation programme (Grant Agreement No. 101017716).

## Author contribution

E. L.-S.: Conceptualization, Methodology, Software, Investigation, Writing - Original Draft, Writing - Review & Editing, Visualization; B. M.: Conceptualization, Methodology, Software, Writing - Review & Editing; E. L.-C.: Methodology, Software, Writing - Review & Editing; R. S.: Methodology, Software, Writing - Review & Editing; R. S.-T.: Conceptualization, Methodology, Visualization; F. W.: Conceptualization, Writing - Review & Editing, Funding; F. B.: Conceptualization, Funding; G. R.: Conceptualization, Methodology, Software, Writing - Review & Editing, Supervision, Funding.

## Ethics statement

This study was approved by the institutional review board of the Assistance Publique Hopitaux de Marseille, and informed written consent was obtained for all patients. All research involving human participants was conducted in accordance with the principles embodied in the Declaration of Helsinki and with local statutory requirements.

## Appendix

### A NMM parameters

The equations of the NMMs used in the model for non-EZ nodes are described in Wendling (2002) [20], and the equations of the NMMs used in EZ nodes, a variation of the non-EZ model including chloride dynamics for seizure initiation, are described in Lopez-Sola (2022) [29].

The parameters of the NMMs used for this study are summarized in Table A1, and are taken from Goodfellow (2016) [12] for the non-EZ models, as explained in the main text, and from Patient 2 of Lopez-Sola (2022) [29]. A key difference of the present work is that the seizure initiation in the EZ NMM is triggered by a change in the external input to the pyramidal population following a step function from 0 to 110 Hz, which leads to the transition to ictal state in the NMM.

**Table A1:**
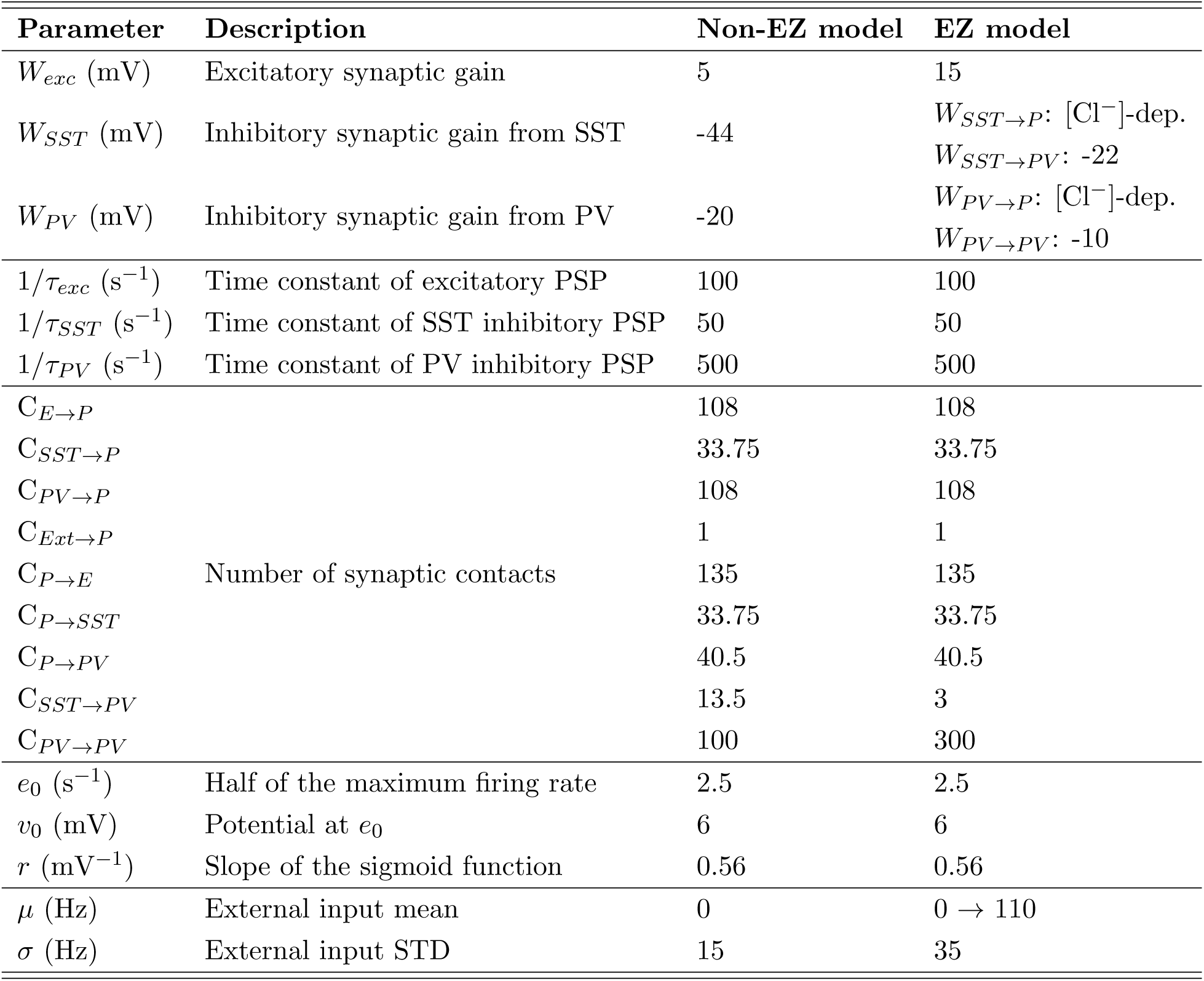
Parameters used in the neural mass models for non-EZ and EZ nodes. The second column provides a brief description of each parameter. The synaptic gains *W_SST →P_* and *W_P V →P_* of the EZ model depend on chloride dynamics.

| Parameter | Description | Non-EZ model | EZ model |
| --- | --- | --- | --- |
| $W_{exc}$ (mV) | Excitatory synaptic gain | 5 | 15 |
| $W_{SST}$ (mV) | Inhibitory synaptic gain from SST | -44 | $W_{SST \rightarrow P}$: $[Cl^-]$-dep.<br>$W_{SST \rightarrow PV}$: -22 |
| $W_{PV}$ (mV) | Inhibitory synaptic gain from PV | -20 | $W_{PV \rightarrow P}$: $[Cl^-]$-dep.<br>$W_{PV \rightarrow PV}$: -10 |
| $1/\tau_{exc}$ ($s^{-1}$) | Time constant of excitatory PSP | 100 | 100 |
| $1/\tau_{SST}$ ($s^{-1}$) | Time constant of SST inhibitory PSP | 50 | 50 |
| $1/\tau_{PV}$ ($s^{-1}$) | Time constant of PV inhibitory PSP | 500 | 500 |
| $C_{E \rightarrow P}$ | Number of synaptic contacts | 108 | 108 |
| $C_{SST \rightarrow P}$ | | 33.75 | 33.75 |
| $C_{PV \rightarrow P}$ | | 108 | 108 |
| $C_{Ext \rightarrow P}$ | | 1 | 1 |
| $C_{P \rightarrow E}$ | | 135 | 135 |
| $C_{P \rightarrow SST}$ | | 33.75 | 33.75 |
| $C_{P \rightarrow PV}$ | | 40.5 | 40.5 |
| $C_{SST \rightarrow PV}$ | | 13.5 | 3 |
| $C_{PV \rightarrow PV}$ | | 100 | 300 |
| $e_0$ ($s^{-1}$) | Half of the maximum firing rate | 2.5 | 2.5 |
| $v_0$ (mV) | Potential at $e_0$ | 6 | 6 |
| $r$ ($mV^{-1}$) | Slope of the sigmoid function | 0.56 | 0.56 |
| $\mu$ (Hz) | External input mean | 0 | $0 \rightarrow 110$ |
| $\sigma$ (Hz) | External input STD | 15 | 35 |

**Table A2:** Parameters, descriptions, and values of the [Cl*^−^*] dynamics of the EZ model. The parameters correspond to Patient 2 in [29].

| Parameter | Description | Value |
| --- | --- | --- |
| $w_0$ (a.u.) | Scaling factor from GABAergic current to mean PSP amplitude | PV→P: 42<br>SST→P: 35 |
| $W_h$ (mV) | Contribution of healthy cells to the average PSP amplitude | PV→P: -290<br>SST→P: -160 |
| $\alpha_{vol}$ (mM·cm<sup>2</sup>/s·μA) | Surface-to-volume and charge-to-concentration factor | PV→P: 0.02<br>SST→P: 0.001 |
| $\alpha_\varphi$ (s·mS/cm<sup>2</sup>) | Scaling factor from GABAergic firing rate to $[\text{Cl}^-]$ currents | 1 |
| $\alpha_{KCC2}$ (mS/cm<sup>2</sup>) | Chloride extrusion rate by KCC2 transporters | SST→P: 1<br>PV→P: 10 |
| $V_m$ (mV) | Average membrane potential | -65 |
| $E_{K^+}$ (mV) | $\text{K}^+$ reversal potential | -85 |
| $[\text{Cl}^-]_{i,\text{base}}^d$ (mM) | Baseline $\text{Cl}^-$ accumulation | PV→P: 10.8<br>SST→P: 8.5 |
| $[\text{Cl}^-]_o$ (mM) | $\text{Cl}^-$ concentration outside the cell | 150 |
| $[\text{HCO}_3^-]_o$ (mM) | $\text{HCO}_3^-$ concentration outside the cell | 25 |
| $[\text{HCO}_3^-]_i$ (mM) | $\text{HCO}_3^-$ concentration inside the cell | 15 |

### B Simulation of SEEG signals

**Figure B1:**
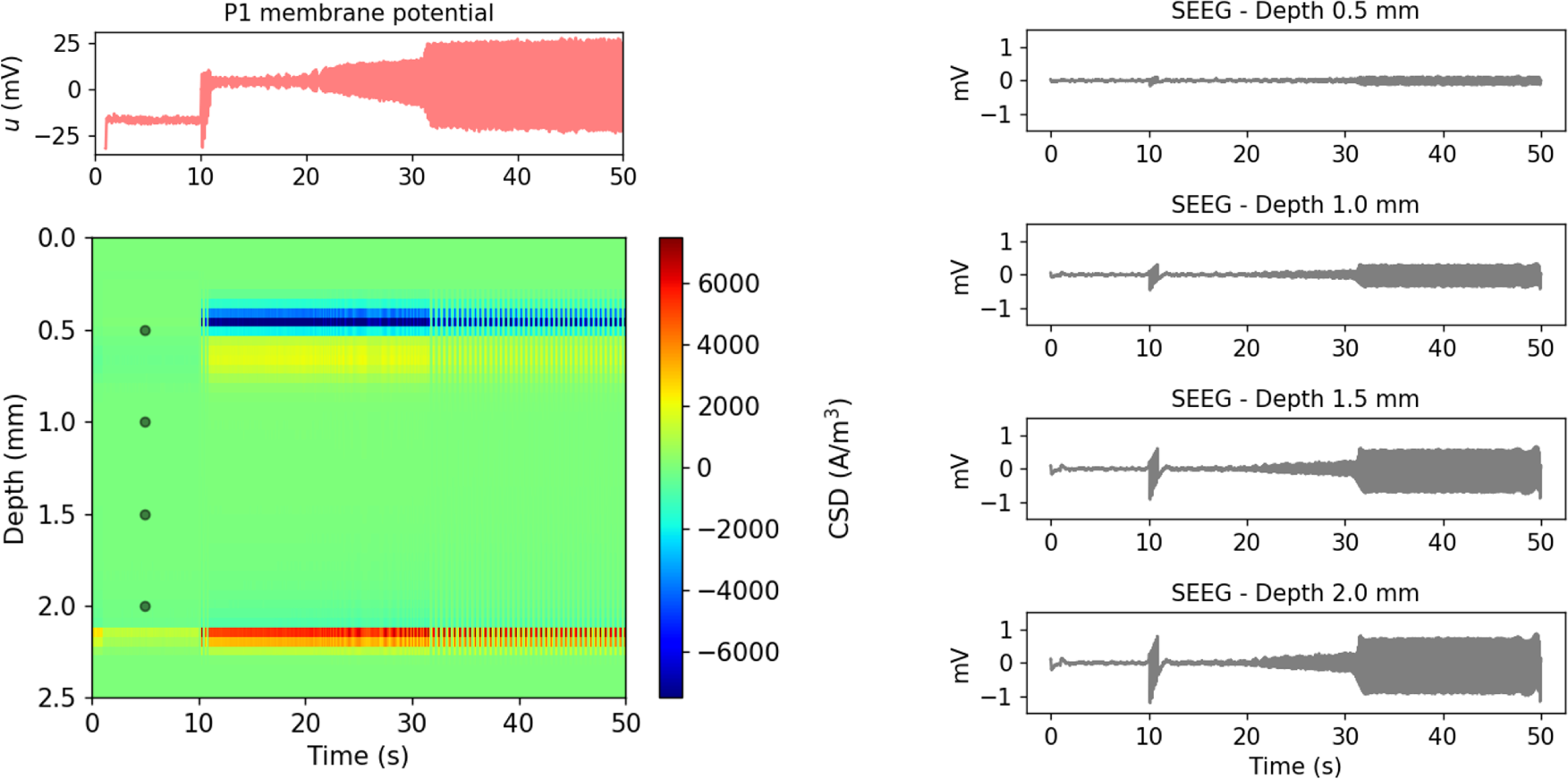
*Left:* Pyramidal population membrane potential (top) and CSD profile (bottom) for the seizure activity of one EZ node. *Right:* Synthetic SEEG signals for different electrode depths (marked by dots in the CSD profile).

**Figure B2:**
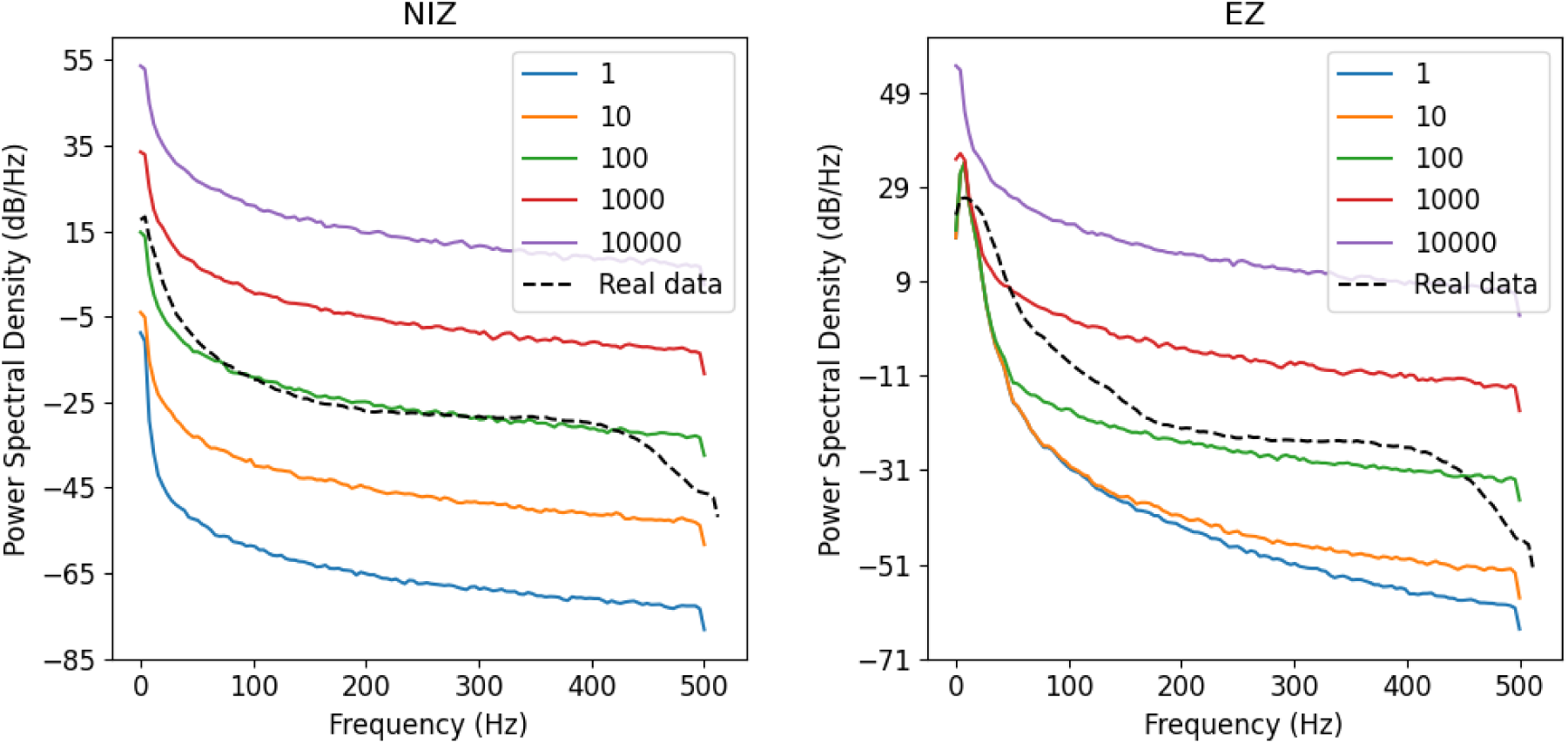
Noise fitting. PSD of empirical and synthetic SEEG data of NIZ nodes (left) and EZ nodes (right) for different scaling factors applied to 1/*f*^2^ noise (in *µ*V). Real data corresponds to representative signals recorded by EZ and NIZ contacts of Patient 2. The optimal value chosen, that best approximates the real data PSD, is 100 *µ*V.

### C Optimization hyperparameters

#### C.1 Regularization term

**Figure C1:**
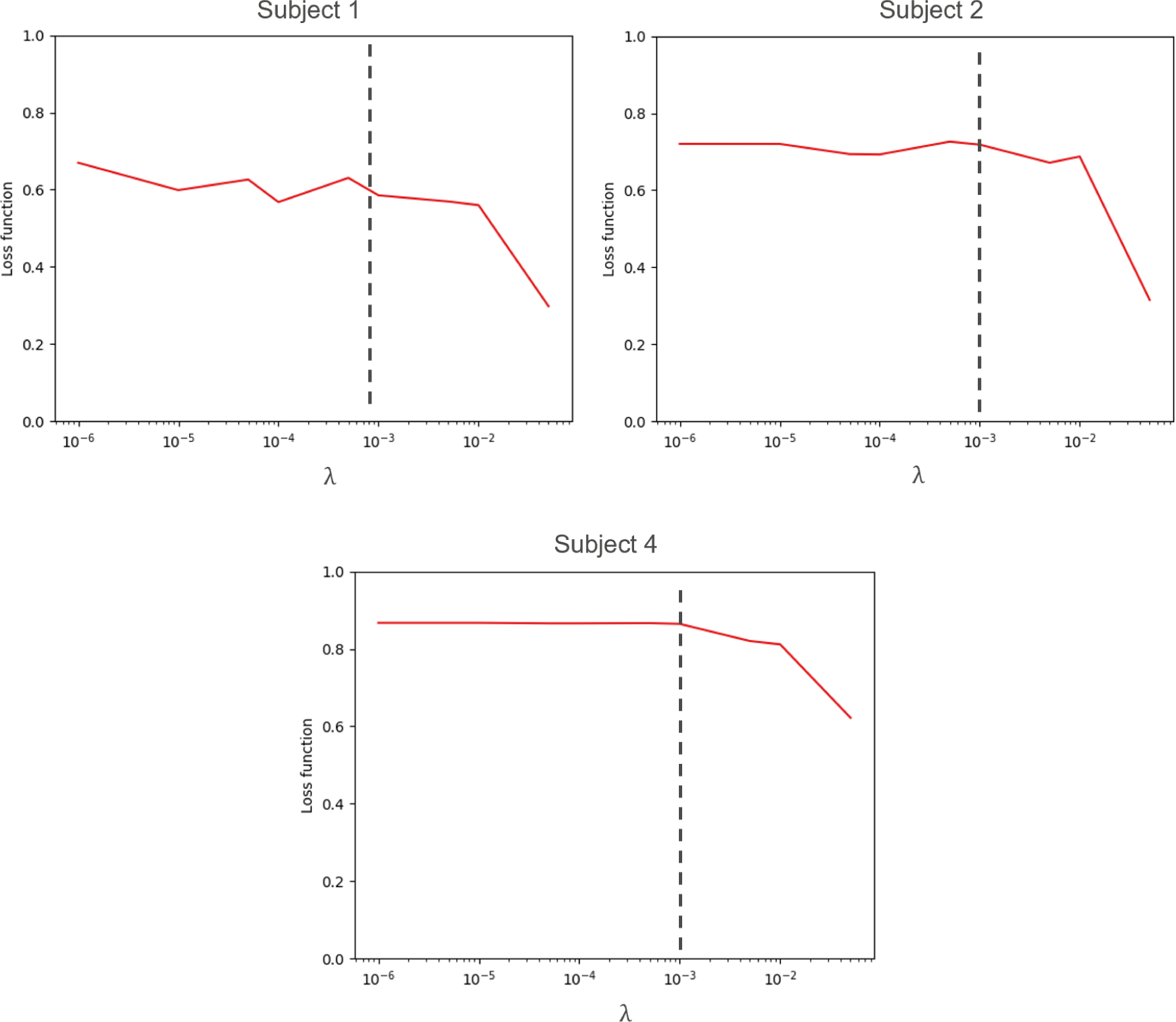
PCC (Loss function) between empirical and synthetic FC values for different values of the regularization term *λ* in three different subjects. The dashed line marks the selected value, where the PCC starts decreasing for most patients.

#### C.2 Hyperparameter tuning

The performance of the algorithm is influenced by four main parameters, the mutation rate, the recombination rate, the population size, and the number of generations. The optimal values for these parameters are problem-dependent and they need to be adjusted on a case-by-case basis. We thus studied the performance of the optimization algorithm for different values of these hyperparameters in terms of PCC between synthetic and empirical FC, convergence, and the variability of the personalized excitability values for one particular patient (Patient 1). Figure C2 shows the evolution of such metrics for different hyperparameter values.

For each hyperparameter studied, we had to fix the other hyperparameters. We used as starting values a crossover rate of 0.4, a mutation rate of (0.3, 0.7), a population multiplier of 20 and a maximum number of generations of 60. For the mutation rate, a range defined by two values is specified because the algorithm employs *dithering*. Dithering helps convergence by randomly changing the mutation constant each generation based on the specified range [62]. The population size is defined by the product of the number of parameters and the population multiplier.

We selected the hyperparameters that resulted in the best PCC between the synthetic and empirical FC, while ensuring a good convergence and a low variability in the personalized excitability values *W_exc_*. The final values selected for the hyperparameters are:

- Population multiplier: 50
- Mutation rate: (0.1, 0.5)
- Crossover rate: 0.4
- Number of generations: 50

**Figure C2:**
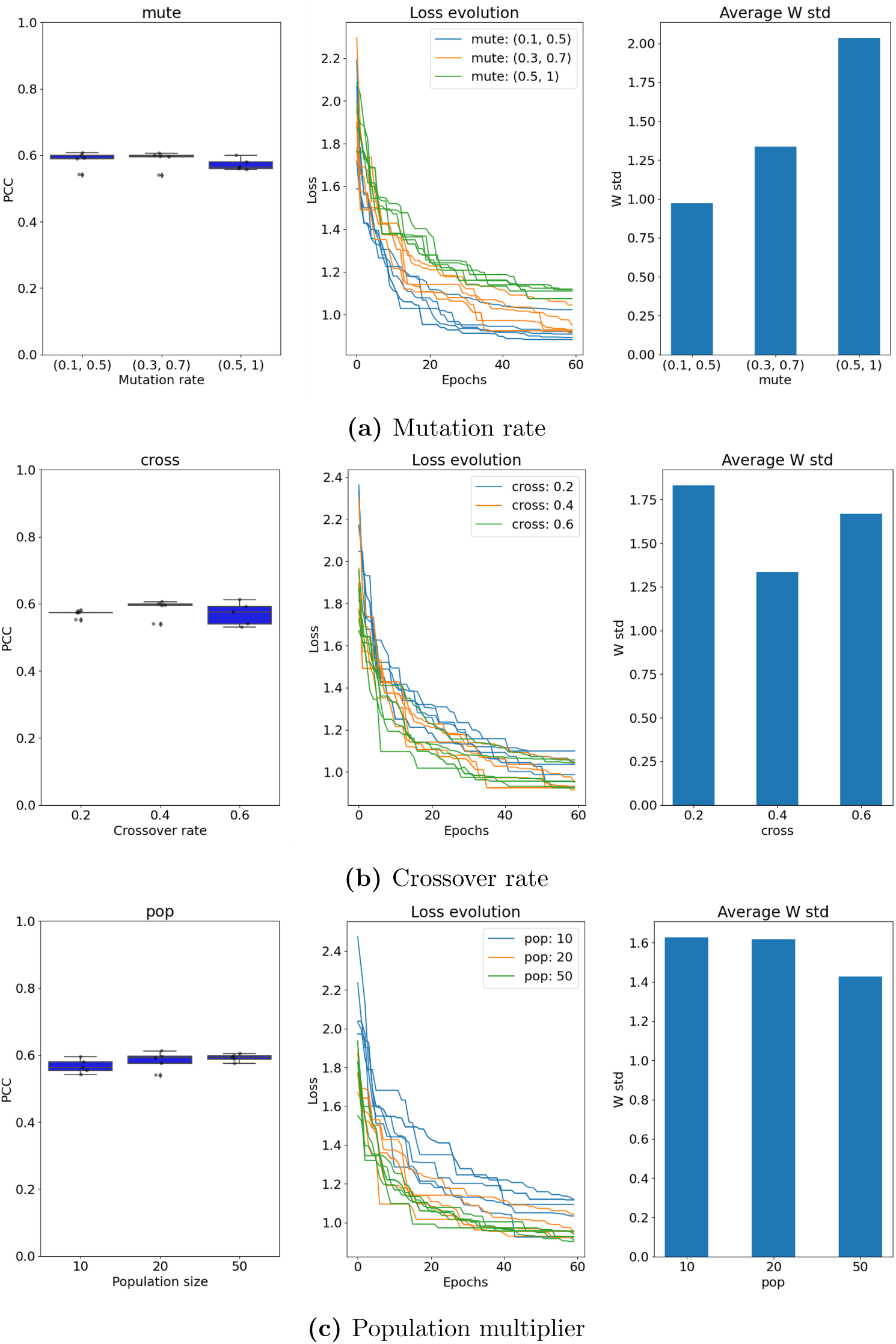
(a) PCC between empirical and synthetic FC (left), evolution of the loss function value (middle) and average standard deviation of the excitability values (right) depending on the **mutation rate** (*mute*). (b) Evaluation of the same metrics depending on the **crossover rate** (*cross*). (c) Evaluation of the same metrics depending on the **population multiplier** *(pop)*.

### D Personalization results (extended)

The values of the personalized parameters for each patient are shown in Figure D1. The comparison of the empirical and synthetic SEEG signals for Patients 4, 5, and 6 is shown in Figure D2 (complementing Figure 5). The presence of seizures in the whole-brain network for each patient is illustrated by Figure D4.

**Figure D1:**
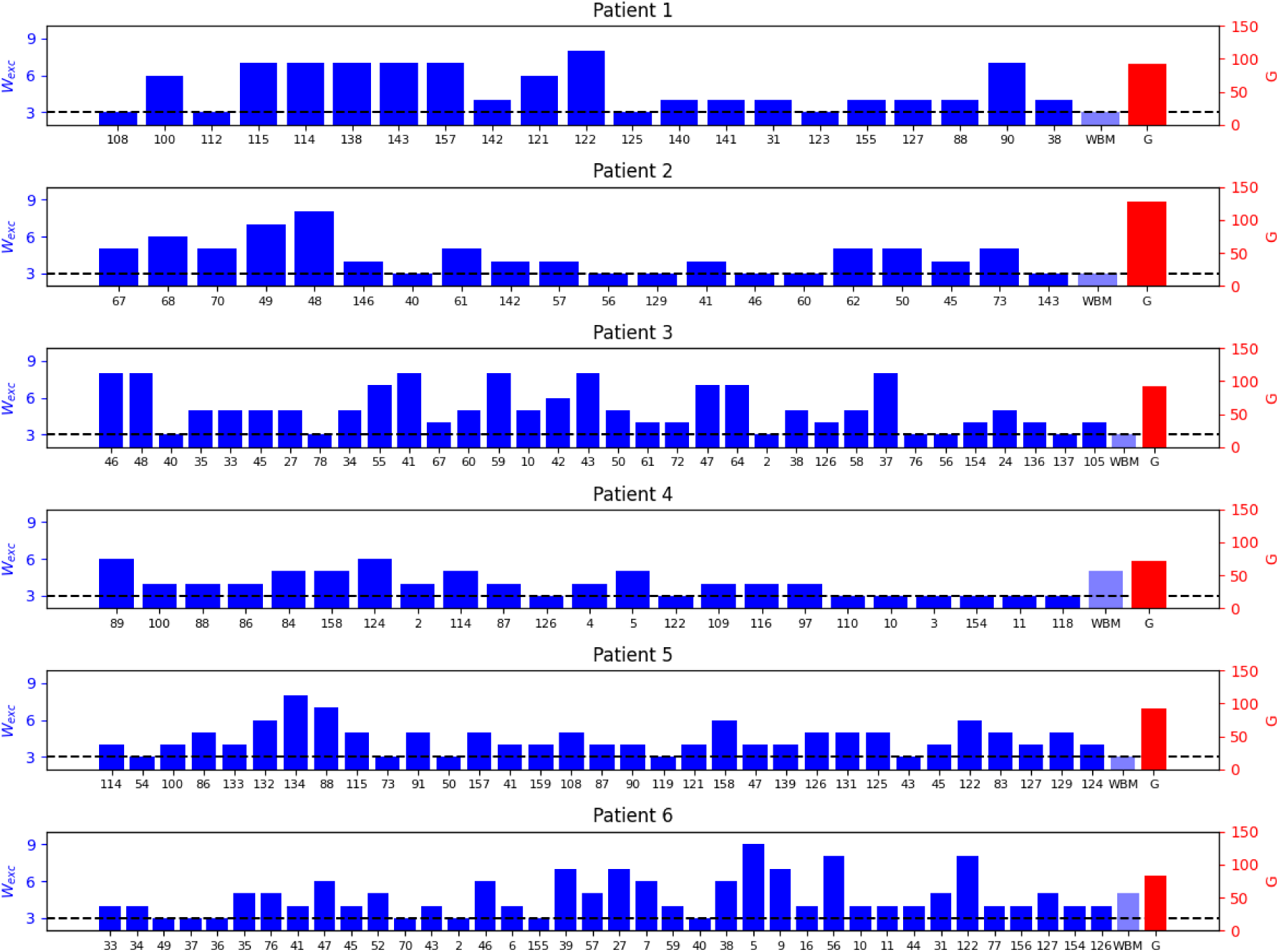
Personalized parameters (G and W*_exc_*) for all the patients in the study. Each panel shows the personalized excitatory synaptic gain 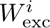 (blue bars, left axis) for each brain region included in the model, along with the global coupling parameter *G* (red bar, right axis). Region labels correspond to VEP parcellation indices. The dashed black line indicates the healthy baseline value of *W*_exc_. WBM refers to the value of excitatory gain for the whole-brain model nodes not sampled by SEEG.

**Figure D2:**
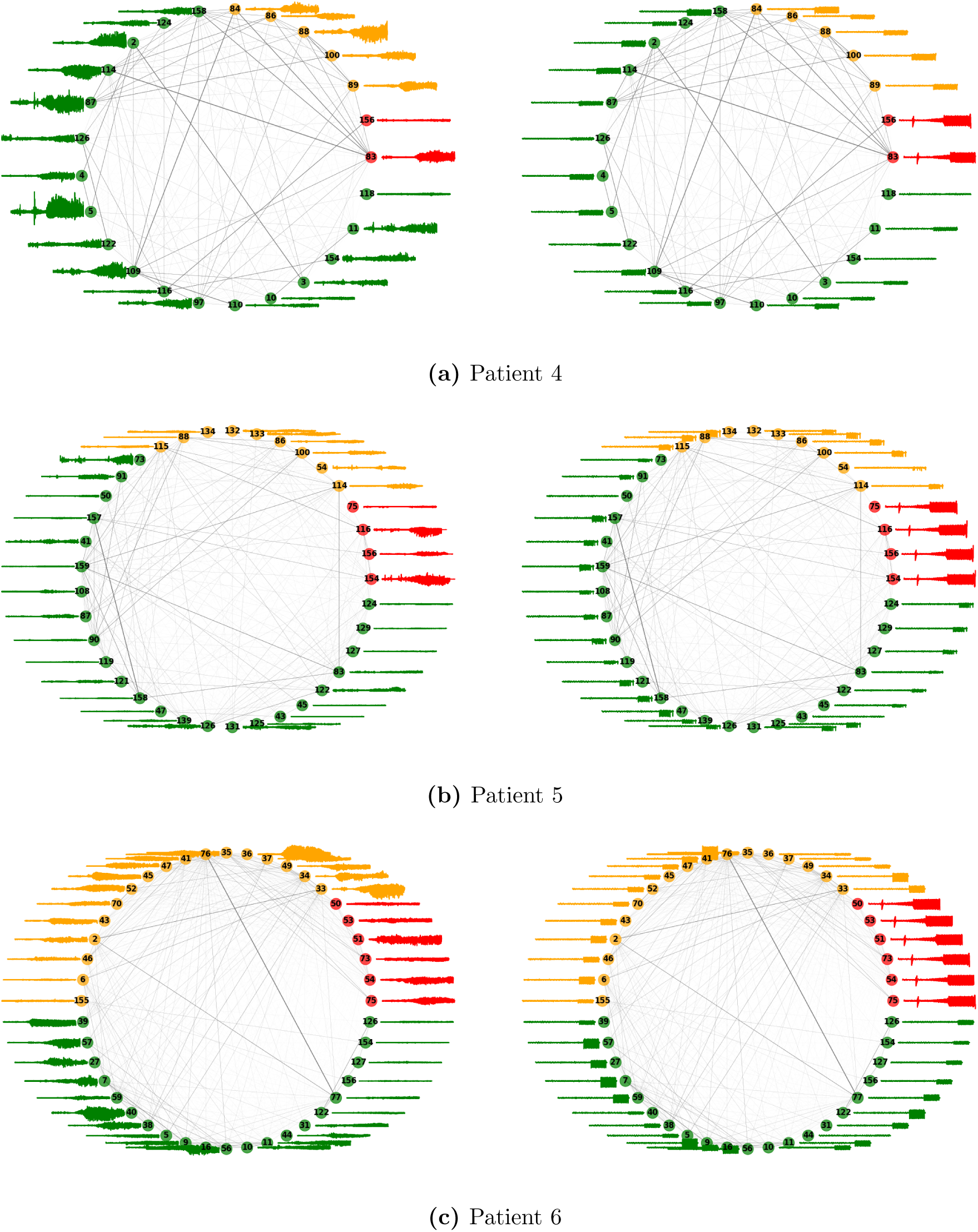
Comparison between empirical (left) and synthetic (right) SEEG signals for Patients 4, 5, and 6. For the empirical data, only one seizure recording is shown. Node numbers represent parcel numbers in the VEP parcellation, and edge width represents the connectivity strength between two nodes. The other three patients are shown in Figure 5.

**Figure D3.**
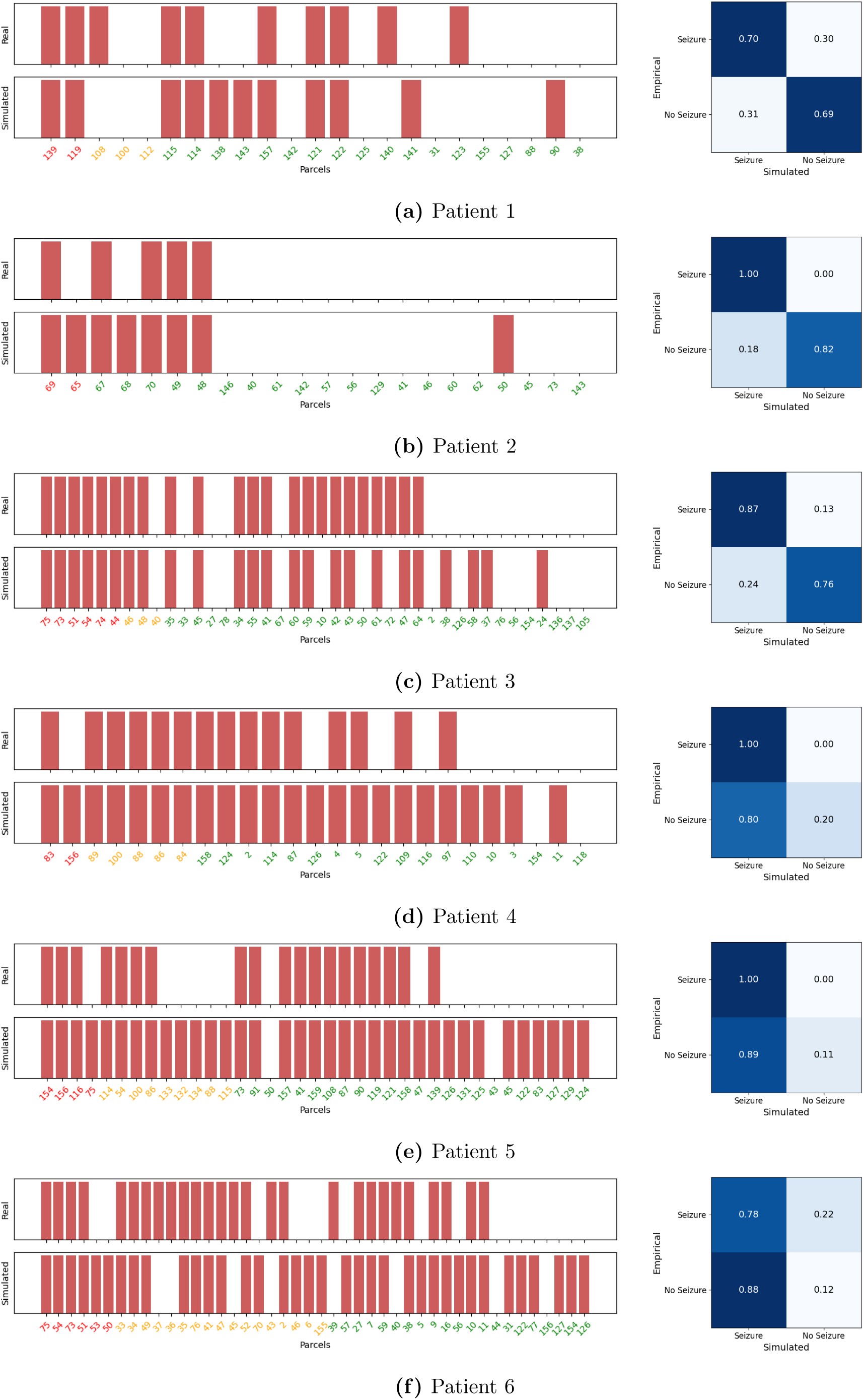
Comparison of empirical and simulated seizure detection. *Left:* Seizure detection across parcels, with red bars indicating regions where seizure activity was detected. Rows correspond to empirical (top) and simulated (bottom) detections. *Right:* Normalized confusion matrices quantifying the agreement between empirical and simulated detections for each patient. Values represent the proportion of parcels in each category (seizure/non-seizure) correctly or incorrectly predicted by the model.

**Figure D4:**
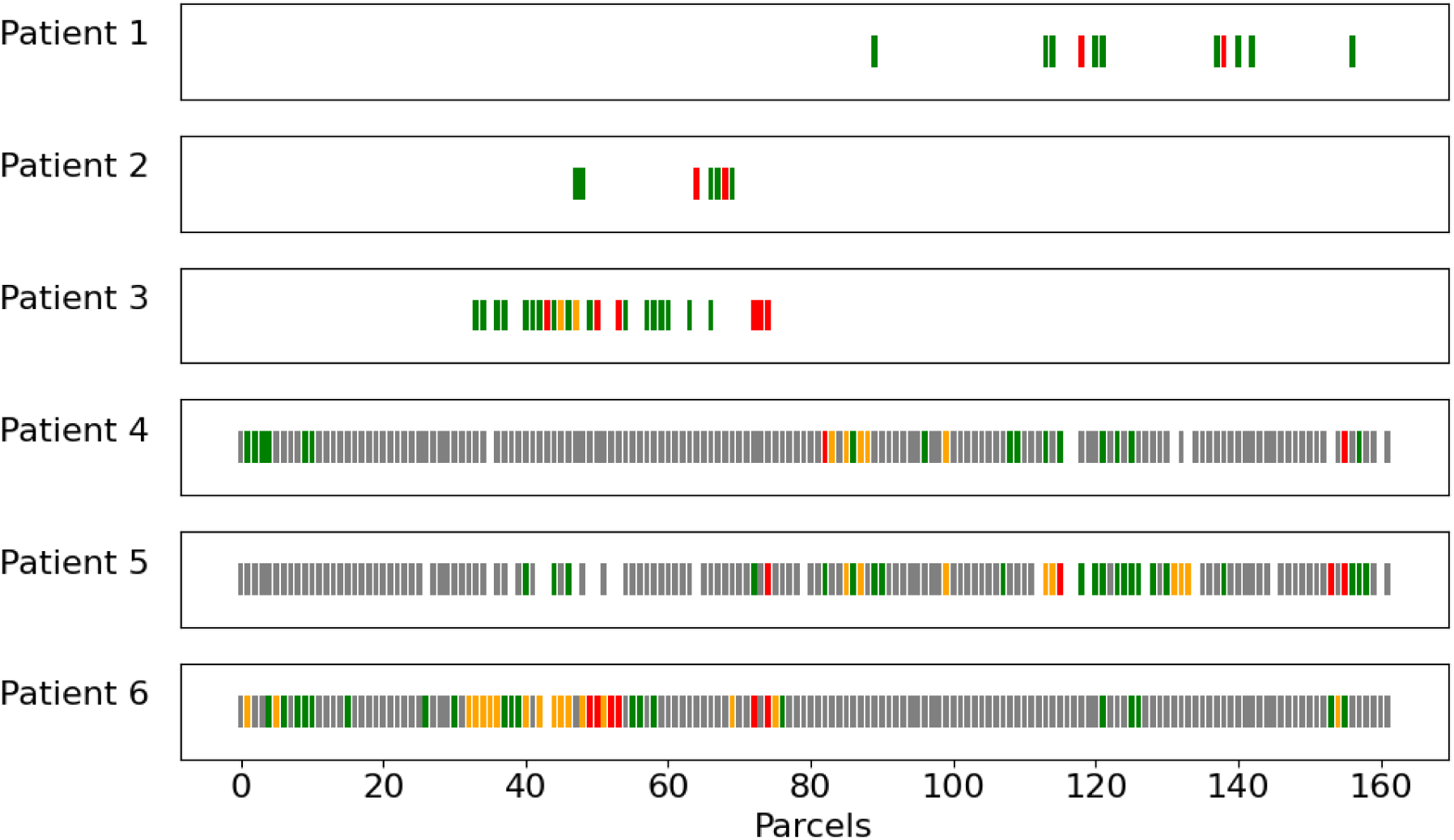
Seizures in the personalized whole-brain model for the six patients in the study. Colors indicate the epileptogenicity of the regions assigned by the clinician: red = EZ, yellow = PZ, green = NIZ, grey = not-sampled by SEEG. Parcel IDs are taken from the VEP parcellation atlas [39].

## References

[1] World Health Organization. Epilepsy: a public health imperative. World Health Organization, 2019. url: https://www.who.int/publications-detail-redirect/epilepsy-a-public-health-imperative.

[2] Patrick Kwan and Martin J Brodie. “Early identification of refractory epilepsy”. In: New England Journal of Medicine 342.5 (2000), pp. 314–319.

[3] Fabrice Wendling. “Computational models of epileptic activity: a bridge between observation and pathophysiological interpretation”. In: Expert review of neurotherapeutics 8.6 (2008), pp. 889–896.

[4] Viktor Jirsa et al. “Personalised virtual brain models in epilepsy”. In: The Lancet Neurology 22.5 (2023), pp. 443–454.

[5] Meysam Hashemi et al. “Principles and Operation of Virtual Brain Twins”. In: IEEE Reviews in Biomedical Engineering (2025), pp. 1–29. doi: 10.1109/RBME.2025.3562951.

[6] Viktor K Jirsa et al. “The virtual epileptic patient: individualized whole-brain models of epilepsy spread”. In: Neuroimage 145 (2017), pp. 377–388.

[7] Meysam Hashemi et al. “The Bayesian Virtual Epileptic Patient: A probabilistic framework designed to infer the spatial map of epileptogenicity in a personalized large-scale brain model of epilepsy spread”. In: NeuroImage 217 (2020), p. 116839.

[8] Moritz Gerster et al. “Patient-specific network connectivity combined with a next generation neural mass model to test clinical hypothesis of seizure propagation”. In: Frontiers in Systems Neuroscience (2021), p. 79.

[9] Marinho A Lopes, Marc Goodfellow, and John R Terry. “A model-based assessment of the seizure onset zone predictive power to inform the epileptogenic zone”. In: Frontiers in computational neuroscience 13 (2019), p. 25.

[10] Viktor Sip et al. “Data-driven method to infer the seizure propagation patterns in an epileptic brain from intracranial electroencephalography”. In: PLoS computational biology 17.2 (2021), e1008689.

[11] Leandro Junges et al. “The role that choice of model plays in predictions for epilepsy surgery”. In: Scientific reports 9.1 (2019), pp. 1–12.

[12] Marc Goodfellow et al. “Estimation of brain network ictogenicity predicts outcome from epilepsy surgery”. In: Scientific reports 6.1 (2016), pp. 1–13.

[13] Nishant Sinha et al. “Predicting neurosurgical outcomes in focal epilepsy patients using computational modelling”. In: Brain 140.2 (2017), pp. 319–332.

[14] Isa Dallmer-Zerbe, Premysl Jiruska, and Jaroslav Hlinka. “Personalized dynamic network models of the human brain as a future tool for planning and optimizing epilepsy therapy”. In: Epilepsia 64.9 (2023), pp. 2221–2238.

[15] Marinho A Lopes et al. “An optimal strategy for epilepsy surgery: Disruption of the rich-club?” In: PLoS computational biology 13.8 (2017), e1005637.

[16] Petroula Laiou et al. “Quantification and selection of ictogenic zones in epilepsy surgery”. In: Frontiers in neurology 10 (2019), p. 1045.

[17] Simona Olmi et al. “Controlling seizure propagation in large-scale brain networks”. In: PLoS computational biology 15.2 (2019), e1006805.

[18] Huifang E Wang et al. “Delineating epileptogenic networks using brain imaging data and personalized modeling in drug-resistant epilepsy”. In: Science Translational Medicine 15.680 (2023), eabp8982.

[19] Sora An et al. “Optimization of surgical intervention outside the epileptogenic zone in the Virtual Epileptic Patient (VEP)”. In: PLoS computational biology 15.6 (2019), e1007051.

[20] F. Wendling et al. “Epileptic fast activity can be explained by a model of impaired GABAergic dendritic inhibition”. eng. In: The European Journal of Neuroscience 15.9 (May 2002), pp. 1499–1508. issn: 0953-816X. doi: 10.1046/j.1460-9568.2002.01985.x.

[21] F Wendling and P Chauvel. “Transition to ictal activity in temporal lobe epilepsy: insights from macroscopic models”. In: Computational neuroscience in epilepsy. Elsevier, 2008, pp. 356–XIV.

[22] Polina Kurbatova et al. “Dynamic changes of depolarizing GABA in a computational model of epileptogenic brain: Insight for Dravet syndrome”. In: Experimental neurology 283 (2016), pp. 57–72.

[23] Elif Koksal-Ersoz et al. “Whole-brain simulation of interictal epileptic discharges for patient-specific interpretation of interictal SEEG data”. In: Neurophysiologie Clinique 54.5 (2024), p. 103005.

[24] Mehmet Alihan Kayabas et al. “Transition to seizure in focal epilepsy: From SEEG phenomenology to underlying mechanisms”. In: Epilepsia 65.12 (2024), pp. 3619–3630.

[25] Emmanouil Giannakakis et al. “Computational modelling of the long-term effects of brain stimulation on the local and global structural connectivity of epileptic patients”. In: Plos one 15.2 (2020), e0221380.

[26] Mäeva Daoud et al. “Stereo-EEG based personalized multichannel transcranial direct current stimulation in drug-resistant epilepsy”. In: Clinical Neurophysiology 137 (2022), pp. 142–151.

[27] Frances Hutchings et al. “Predicting surgery targets in temporal lobe epilepsy through structural connectome based simulations”. In: PLoS computational biology 11.12 (2015), e1004642.

[28] George Petkov et al. “A critical role for network structure in seizure onset: a computational modeling approach”. In: Frontiers in neurology 5 (2014), p. 261.

[29] Edmundo Lopez-Sola et al. “A personalizable autonomous neural mass model of epileptic seizures”. In: Journal of Neural Engineering 19.5 (2022), p. 055002.

[30] Roser Sanchez-Todo et al. “A physical neural mass model framework for the analysis of oscillatory generators from laminar electrophysiological recordings”. In: NeuroImage 270 (2023), p. 119938.

[31] Borja Mercadal et al. “Towards a mesoscale physical modeling framework for stereotactic-EEG recordings”. In: Journal of Neural Engineering 20.1 (2023), p. 016005.

[32] Andrew J Trevelyan et al. “On brain stimulation in epilepsy”. In: Brain (2025), awae385.

[33] Philippe Ryvlin et al. “Neuromodulation in epilepsy: state-of-the-art approved therapies”. In: The Lancet Neurology 20.12 (2021), pp. 1038–1047.

[34] Sara Simula et al. “Transcranial current stimulation in epilepsy: a systematic review of the fundamental and clinical aspects”. In: Frontiers in Neuroscience 16 (2022), p. 909421.

[35] Jean-Pascal Lefaucheur and Fabrice Wendling. “Mechanisms of action of tDCS: A brief and practical overview”. In: Neurophysiologie Clinique 49.4 (2019), pp. 269–275.

[36] Harper Lee Kaye et al. “Personalized, multisession, multichannel transcranial direct current stimulation in medication-refractory focal epilepsy: an open-label study”. In: Journal of Clinical Neurophysiology 40.1 (2023), pp. 53–62.

[37] Giulio Ruffini et al. “Targeting brain networks with multichannel transcranial current stimulation (tCS)”. In: Current Opinion in Biomedical Engineering 8 (2018), pp. 70–77.

[38] Jane De Tisi et al. “The long-term outcome of adult epilepsy surgery, patterns of seizure remission, and relapse: a cohort study”. In: The Lancet 378.9800 (2011), pp. 1388–1395.

[39] Huifang E Wang et al. “VEP atlas: An anatomic and functional human brain atlas dedicated to epilepsy patients”. In: Journal of neuroscience methods 348 (2021), p. 108983.

[40] Sara Simula et al. “Impact of transcranial electrical stimulation on simultaneous stereoelectroencephalography recordings: A randomized sham-controlled study”. In: Clinical Neurophysiology 166 (2024), pp. 211–222.

[41] Fabrice Bartolomei et al. “Defining epileptogenic networks: contribution of SEEG and signal analysis”. In: Epilepsia 58.7 (2017), pp. 1131–1147.

[42] S Medina Villalon et al. “EpiTools, A software suite for presurgical brain mapping in epilepsy: Intracerebral EEG J. Neurosci”. In: Methods (2018), pp. 3037–15.

[43] Julia Makhalova et al. “Virtual epileptic patient brain modeling: Relationships with seizure onset and surgical outcome”. In: Epilepsia 63.8 (2022), pp. 1942–1955.

[44] B. Fischl et al. “Automatically parcellating the human cerebral cortex”. In: Cerebral cortex 14.1 (2004), pp. 11–22. doi: 10.1093/cercor/bhg087.

[45] J-Donald Tournier et al. “MRtrix3: A fast, flexible and open software framework for medical image processing and visualisation”. In: Neuroimage 202 (2019), pp. 116–137.

[46] Lacoma Ayoubian, H Lacoma, and J Gotman. “Automatic seizure detection in SEEG using high frequency activities in wavelet domain”. In: Medical engineering & physics 35.3 (2013), pp. 319–328.

[47] Pieter Van Mierlo et al. “Ictal-onset localization through connectivity analysis of intracranial EEG signals in patients with refractory epilepsy”. In: Epilepsia 54.8 (2013), pp. 1409–1418.

[48] Stanislas Lagarde et al. “Interictal functional connectivity in focal refractory epilepsies investigated by intracranial EEG”. In: Brain connectivity 12.10 (2022), pp. 850–869.

[49] Fabrice Wendling et al. “From EEG signals to brain connectivity: a model-based evaluation of interdependence measures”. In: Journal of neuroscience methods 183.1 (2009), pp. 9–18.

[50] Stanislas Lagarde et al. “The repertoire of seizure onset patterns in human focal epilepsies: determinants and prognostic values”. In: Epilepsia 60.1 (2019), pp. 85–95.

[51] Fabrice Wendling et al. “From intracerebral EEG signals to brain connectivity: identification of epileptogenic networks in partial epilepsy”. In: Frontiers in systems neuroscience 4 (2010), p. 154.

[52] Gustavo Deco et al. “Resting-state functional connectivity emerges from structurally and dynamically shaped slow linear fluctuations”. In: Journal of Neuroscience 33.27 (2013), pp. 11239–11252.

[53] Maxim Bazhenov et al. “Cellular and network mechanisms of electrographic seizures”. In: Drug Discovery Today: Disease Models 5.1 (2008), pp. 45–57.

[54] Ben H Jansen and Vincent G Rit. “Electroencephalogram and visual evoked potential generation in a mathematical model of coupled cortical columns”. In: Biological cybernetics 73.4 (1995), pp. 357–366.

[55] G. Ruffini et al. “P118 A Biophysically Realistic Laminar Neural Mass Modeling Framework for Transcranial Current Stimulation”. In: Clinical Neurophysiology 131.4 (Apr. 1, 2020), e78–e79. issn: 1388-2457. doi: 10.1016/j.clinph.2019.12.229. url: https://www.sciencedirect.com/science/article/pii/S1388245719315950.

[56] Roser Sanchez-Todo et al. “Fast Interneuron Dysfunction in Laminar Neural Mass Model Reproduces Alzheimer’s Oscillatory Biomarkers”. In: biorxiv (2025), pp. 2025–03.

[57] Gÿorgy Buzśaki, Costas A Anastassiou, and Christof Koch. “The origin of extracellular fields and currents—EEG, ECoG, LFP and spikes”. In: Nature reviews neuroscience 13.6 (2012), pp. 407–420.

[58] Paul L Nunez, Ramesh Srinivasan, et al. Electric fields of the brain: the neurophysics of EEG. Oxford University Press, USA, 2006.

[59] F Wendling et al. “Brain (hyper) excitability revealed by optimal electrical stimulation of GABAergic interneurons”. In: Brain Stimulation 9.6 (2016), pp. 919–932.

[60] Fabrice Wendling et al. “Multiscale neuro-inspired models for interpretation of EEG signals in patients with epilepsy”. In: Clinical Neurophysiology 161 (2024), pp. 198–210.

[61] Rainer Storn and Kenneth Price. “Differential Evolution – A Simple and Efficient Heuristic for global Optimization over Continuous Spaces”. In: Journal of Global Optimization 11.4 (1997), pp. 341–359. doi: 10.1023/a:1008202821328.

[62] Scipy differential evolution documentation. https://docs.scipy.org/doc/scipy/reference/generated/scipy.optimize.differential_evolution.html. Accessed: 2023-11-27.

[63] Philip N Patsalos et al. “Antiepileptic drugs—best practice guidelines for therapeutic drug monitoring: a position paper by the subcommission on therapeutic drug monitoring, ILAE Commission on Therapeutic Strategies”. In: Epilepsia 49.7 (2008), pp. 1239–1276.

[64] Shery Jacob and Anroop B Nair. “An updated overview on therapeutic drug monitoring of recent antiepileptic drugs”. In: Drugs in R&D 16 (2016), pp. 303–316.

[65] Max O. Krucoff et al. “Operative Technique and Lessons Learned From Surgical Implantation of the NeuroPace Responsive Neurostimulation® System in 57 Consecutive Patients”. In: Operative Neurosurgery (Hagerstown, Md.) 20.2 (Jan. 13, 2021), E98–E109. issn: 2332-4260. doi: 10.1093/ons/opaa300. PMID: 33074294.

[66] Carl E Stafstrom. “Distinct Mechanisms Mediate Interictal and Pre-Ictal Discharges in Human Temporal Lobe Epilepsy: Interictal and Pre-Ictal Discharges”. In: Epilepsy Currents 11.6 (2011), pp. 200–202.

[67] Olesya Grinenko et al. “A fingerprint of the epileptogenic zone in human epilepsies”. In: Brain 141.1 (2018), pp. 117–131.

[68] Borja Mercadal et al. “tDCS montage optimization for the treatment of epilepsy using Neurotwins”. In: Neuromodec Journal 4.2 (2023). doi: 10.31641/nmj-RCWA5184.

[69] Borja Mercadal et al. “tDCS montage optimization for the treatment of epilepsy using personalized whole-brain models”. In prep. 2025.

[70] Yves Denoyer et al. “Modelling acute and lasting effects of tDCS on epileptic activity”. In: Journal of Computational Neuroscience 48.2 (2020), pp. 161–176.

